# Architecture and dynamics of a novel desmosome-endoplasmic reticulum organelle

**DOI:** 10.1101/2022.07.07.499185

**Authors:** Navaneetha Krishnan Bharathan, William Giang, Jesse S. Aaron, Satya Khuon, Teng-Leong Chew, Stephan Preibisch, Eric T. Trautman, Larissa Heinrich, John Bogovic, Davis Bennett, David Ackerman, Woohyun Park, Alyson Petruncio, Aubrey V. Weigel, Stephan Saalfeld, COSEM Project Team, A. Wayne Vogl, Sara N. Stahley, Andrew P. Kowalczyk

## Abstract

The endoplasmic reticulum (ER) forms a dynamic network that contacts other cellular membranes to regulate stress responses, calcium signaling, and lipid transfer. Using high-resolution volume electron microscopy, we find that the ER forms a previously unknown association with keratin intermediate filaments and desmosomal cell-cell junctions. Peripheral ER assembles into mirror image-like arrangements at desmosomes and exhibits nanometer proximity to keratin filaments and the desmosome cytoplasmic plaque. ER tubules exhibit stable associations with desmosomes, and perturbation of desmosomes or keratin filaments alters ER organization and mobility. These findings indicate that desmosomes and the keratin cytoskeleton pattern the distribution of the ER network. Overall, this study reveals a previously unknown subcellular architecture defined by the structural integration of ER tubules with an epithelial intercellular junction.

**One-Sentence Summary:** The desmosome adhesive junction regulates the organization and dynamics of the endoplasmic reticulum network.

## Main Text

The endoplasmic reticulum (ER) is the largest and perhaps most architecturally complex membranous organelle in eukaryotic cells (*1*). The ER participates in protein biosynthesis and turnover, organelle biogenesis, the transfer of lipids between membranous compartments, and regulation of calcium homeostasis (*2, 3*). These functions are often conducted at membrane contact sites between the ER and other organelles including endosomes, mitochondria, and the plasma membrane (PM) (*1, 2*). For these reasons, it is of particular interest to understand how the spatiotemporal behavior of ER membranes is regulated. The morphology and dynamics of the ER have been studied predominantly in individual mammalian cells, including fibroblasts and COS-7 (*4, 5*). However, little is known about how ER tubule organization and dynamics are coordinated between cells that form extensive cell-cell contacts.

Using cryo-fixation approaches to preserve near-native cell structure, in combination with high-resolution volume electron microscopy (*6*), we find that desmosomes, an adhesive cell-cell junction coupled to intermediate filament networks, organize the subcellular distribution and dynamics of ER tubules. Peripheral ER tubules follow keratin filament bundles to desmosomes where they are organized and stabilized in symmetrical arrangements at opposing cell-cell contacts. Disruption of desmosomes or expression of disease-causing keratin mutants alters ER morphology and dynamics. These findings reveal the architecture of a symmetrical organelle assembly at intercellular contacts that comprises desmosomes, intermediate filaments, and the ER. This newly identified role for desmosomes in regulating ER morphology and dynamics provides insights into both fundamental organelle biology and into human disease states resulting from desmosome or keratin dysfunction.

## Results

### The endoplasmic reticulum associates with the desmosome

To assess the organization of ER in epithelial cells, we used previously described A431 cell lines expressing desmoplakin-EGFP to visualize desmosomal cell-cell junctions (*7*), and stably transduced these cells with mApple-VAPB to visualize ER (*8, 9*). Spinning disk confocal microscopy revealed that peripheral ER tubules associated with desmoplakin puncta in symmetrical arrangements on both sides of the desmosomal cell-cell contact (Fig. S1A). ER tubule association with desmosomes was confirmed in multiple other epithelial cell types (Fig. S1B, C). Transmission electron microscopy of parental A431 cells revealed that peripheral ER tubules extend toward the electron-dense desmosome plaque, often exhibiting a symmetrical configuration on either side of the desmosomal junction (Fig. S1D-G).

We determined the nanoscale three-dimensional architecture of ER-desmosome associations using a cryo-structured illumination microscopy (CryoSIM) and focused ion beam scanning electron microscopy (FIB-SEM) workflow (*6*) (Fig. S2). The cytoplasmic zone of the desmosome comprises an electron-dense outer plaque adjacent to the plasma membrane and a less electron-dense inner plaque that functions as an attachment zone for keratin intermediate filaments (*10, 11*). FIB-SEM imaging and 3D reconstructions revealed that peripheral ER tubules were in close contact with desmosomal plaques (Fig. 1). In many instances, these tubules then branched around the desmosome inner dense plaque and traveled toward the plasma membrane at the edges of the outer dense plaque (Fig. 1A, B; Movie S1). ER tubules and keratin filaments were co-organized in mirror image-like arrangements proximal to desmosomal cell-cell contacts (Fig. 1C-K). As shown in Fig. S3 and Movie S2, ER tubules were also observed in the space between the keratin filament attachment zone (inner dense plaque) and the outer dense plaque of the desmosome adjacent to the plasma membrane (arrows, Fig. S3G-I). From a total of 32 desmosomes across the FIB-SEM datasets, ER tubules were found to be within 30nm of the outer dense plaques of 16 desmosomes (50%). In four desmosomes, ER tubules were found to be in contact with the outer dense plaque. These data indicate that peripheral ER tubules are a previously unrecognized component of the desmosomal adhesive complex.

**Fig. 1.**
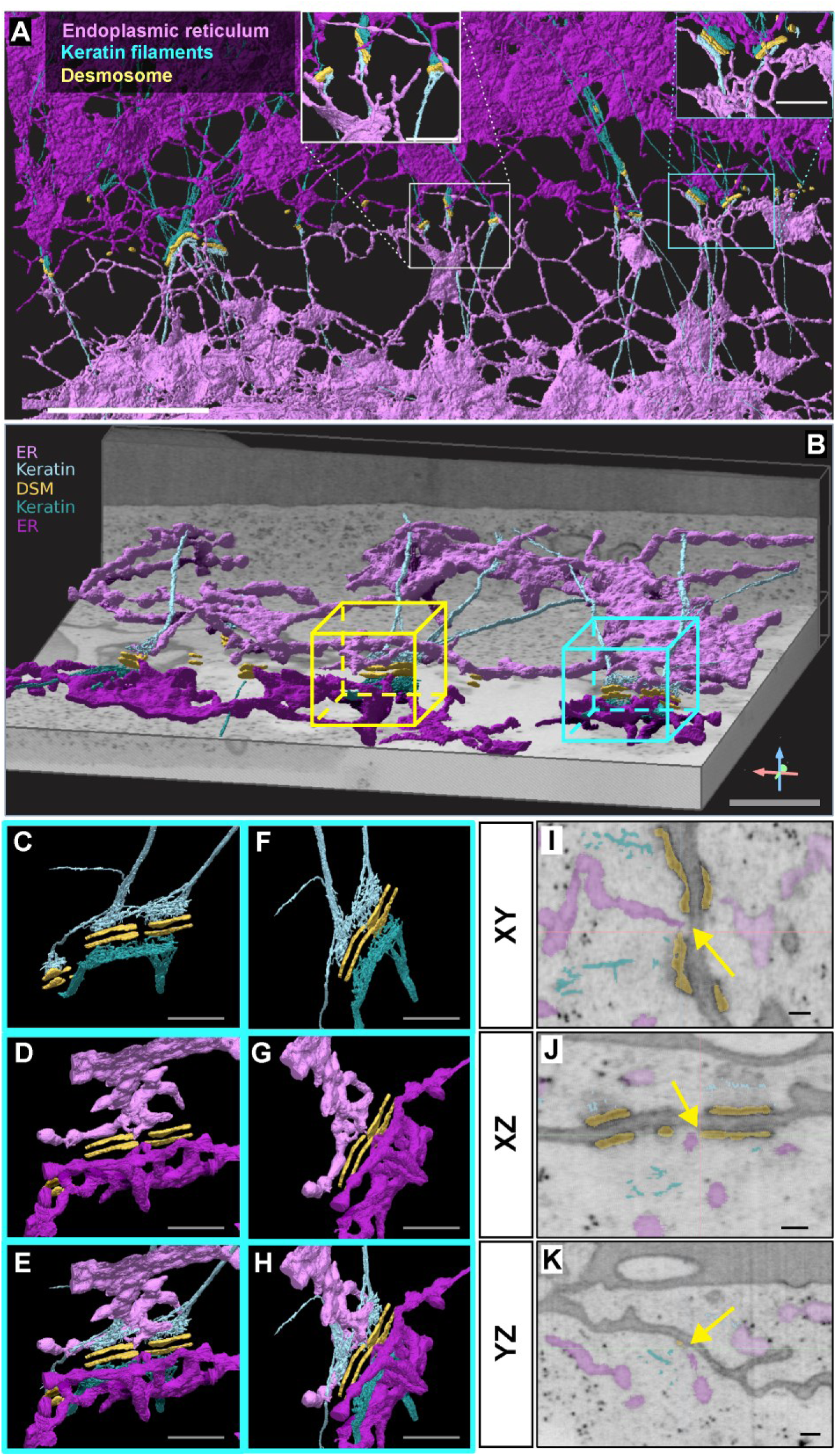
FIB-SEM reveals ER-desmosome associations. (**A**) FIB-SEM segmentations of a cell-cell contact in A431 cells acquired at 8nm isotropic voxel size showing desmosome outer dense plaque (orange), keratin filaments (teal) and ER (magenta). Different shades of teal and magenta distinguish keratin and ER in adjacent cells. White and blue boxes indicate regions with magnified views shown in insets. (**B**) FIB-SEM segmentations of a cell-cell contact in A431 cells acquired at 4nm isotropic voxel size. Teal box indicates region with magnified segmentations shown in C-H. (**C-H**) Rotated views of a desmosome outer dense plaque (orange) showing symmetrical organization of teal keratin filaments (**C, F**) and magenta ER (**D, G**). Panels **E** and **H** show rotated views of desmosome outer dense plaque, keratin filaments and ER. (**I-K**) Orthoslices in XY (**I**), XZ (**J**), and YZ (**K**) with desmosome, ER, and keratin segmentations. Yellow arrows point to ER tubules proximal to desmosome outer dense plaque. Scale bar = 5µm (A), 1µm (B), 500nm (C-H),100nm (I-K).

### The endoplasmic reticulum associates with keratin filaments

The keratin intermediate filament cytoskeleton anchors to the desmosome inner dense plaque, leading to the formation of a symmetrical adhesion structure (*11*). Since peripheral ER tubules were observed on either side of the desmosome, we investigated ER tubule localization relative to keratin filament bundles proximal and distal to desmosomal cell contacts. FIB-SEM datasets at 4×4×4nm^3^ voxel size revealed that peripheral ER tubules intertwined around keratin filaments that approached the desmosome plaque (Fig. 2). Keratin filament bundles were often fully enveloped by ER membrane (Fig. 2A-C and Movie S3). Furthermore, ER tubules often ran parallel to keratin filaments proximal to the desmosome plaque (Fig. 2G-I). We observed frequent instances where ER tubules contacted keratin filament bundles (Fig. 2D-F; J-L). Keratin filaments made similar close contacts with sheet-like/planar ER structures located distal to cell borders (not shown). These observations demonstrate an intimate physical association between the ER membrane and the intermediate filament cytoskeleton. Overall, our FIB-SEM analysis revealed a previously unknown symmetrical organelle assembly comprising ER tubules, keratin filaments, and the desmosome.

**Fig. 2.**
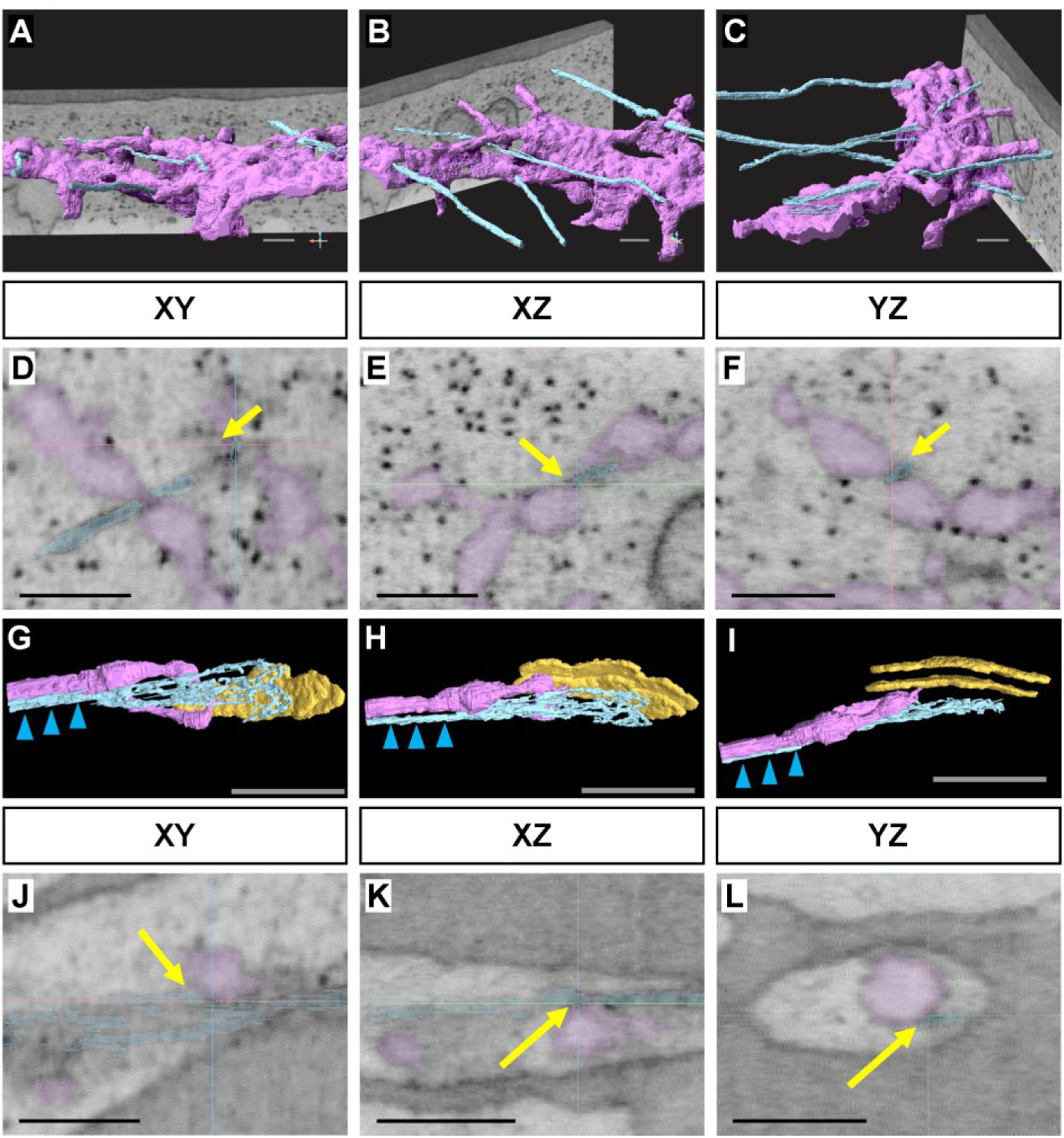
FIB-SEM reveals ER-keratin filament associations in A431 cells. (**A-C**) Rotated views of the same ROI showing keratin filaments (teal) penetrating the ER network (magenta). (**D-F**) Orthoslices of region shown in A-C in XY (**D**), XZ (**E**), and YZ (**F**) with ER and keratin segmentations. (**G-I**) Rotated views of the same ROI showing keratin filaments (teal) proximal to a peripheral ER tubule (magenta) as they approach a desmosome (orange). (**J-L**) Orthoslices of region shown in G-I in XY (**G**), XZ (**H**), and YZ (**I**) with desmosome, ER, and keratin segmentations. Blue arrowheads indicate keratin and ER tubule parallel to each other. Yellow arrows point to contact between ER and keratin at the same point in all 3 views (D-F and J-L). Scale bar = 250nm (A-F; J-L), 500nm (G-I).

### Peripheral ER tubules stably anchor to desmosomes

The ER tubule network is highly dynamic but also forms stable tethers with various membraneless organelles, such as P-bodies (*12*), and membrane-bound organelles, including mitochondria (*13*) and endosomes (*1, 14, 15*). Further, ER tubule dynamics vary depending on tubule associations (*5, 16*). To determine if ER associations with desmosomes observed by FIB-SEM influence ER dynamics, we utilized spinning disk confocal microscopy of living A431 cell lines stably expressing desmoplakin-EGFP and mApple-VAPB to visualize the dynamic relationships between ER and desmosomes. Cells were imaged at an interval of 5 seconds for a duration of 2 minutes (25 timepoints). ER tubules were present at the cell periphery and closely associated with virtually all desmosomes (Fig. 3A; Movie S4). Kymographs revealed that ER tubules associated with desmosomes were highly stable (Fig. 3B). Quantification of ER-desmosome associations revealed that ∼77% of desmosomes (n=154 desmosomes total) were in contact with ER tubules for the entirety of a 2-minute time course, while the remaining 23% of desmosomes made frequent but transient contacts with ER tubules (Fig. 3C; Fig. S4).

**Fig. 3.**
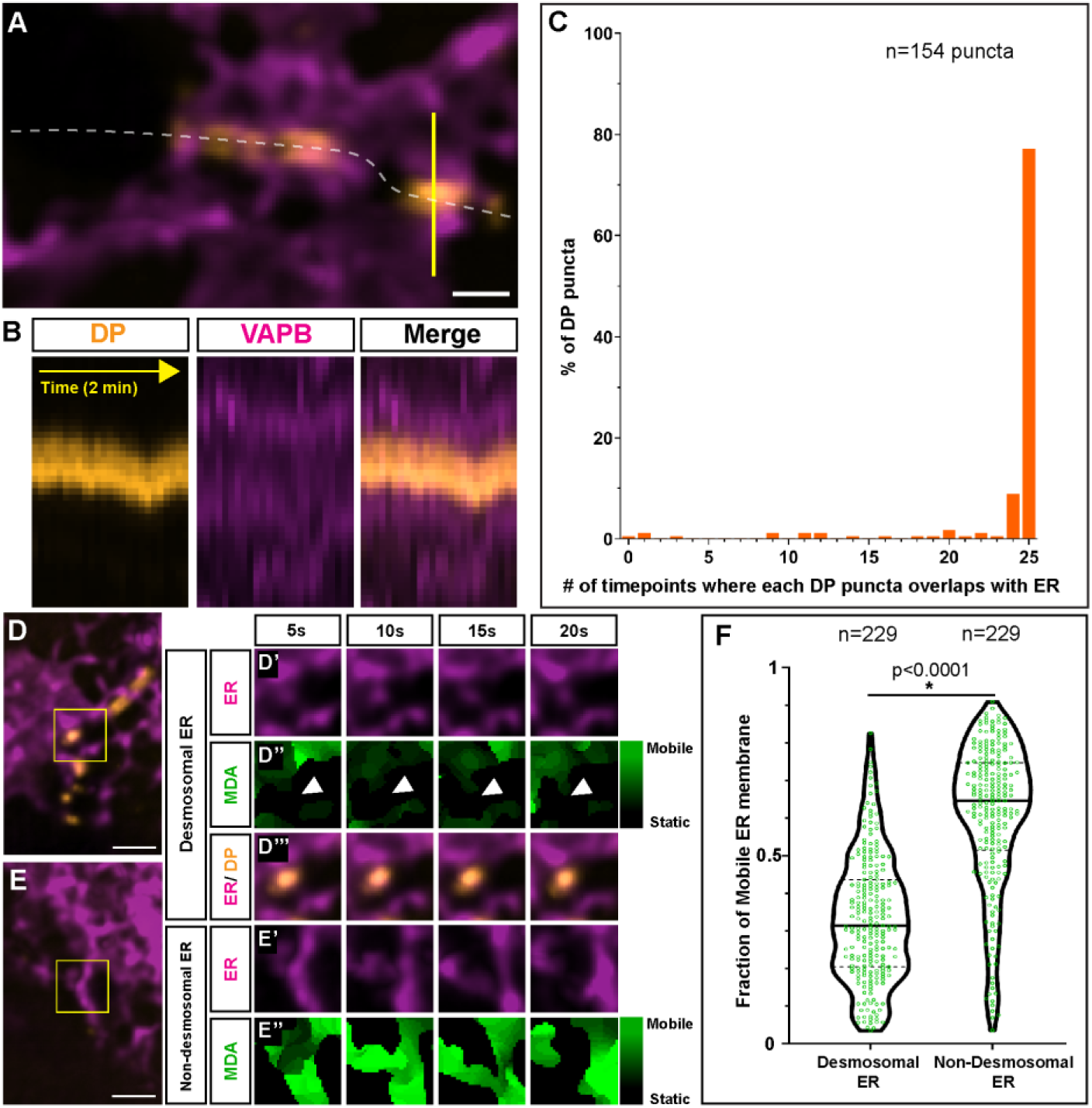
Desmosomes anchor ER tubules and stabilize ER membrane. (**A**) Snapshot of a pair of A431 cells expressing Desmoplakin-EGFP (orange, desmosome marker) and mApple-VAPB (magenta, ER marker) showing ER tubules anchored on either side of desmoplakin puncta. Cell border shown as dashed white line. Solid yellow line indicates position of kymograph in B. (**B**) Kymograph of yellow line in A revealing stable ER-DP contacts over a 2-minute time course. (**C**) Histogram shows percentage of DP puncta that make contacts with ER for *t* timepoints over a 25 timepoint duration. (**D-E**) Yellow boxes highlight regions analyzed by Membrane Displacement Analysis (MDA). (D’-D’’’) ER (magenta), MDA-generated ER movement (green), ER at desmosome contacts (orange). White arrowheads in D’’ depict location of DP puncta. (E’-E’’) ER (magenta), MDA-generated ER movement (green) at non-desmosomal regions. Bright green pixels in D’’ and E’’ depict ER fraction that is mobile between time points. Note more bright green pixels in E’’ vs. D’’, indicating more mobile ER. (**F**) Violin plot depicting fraction of mobile ER in desmosomal vs. non-desmosomal (cytoplasmic) regions. Horizontal black lines in violin plots represent medians (Mann-Whitney, p<0.0001) and *n* indicates number of ROI analyzed. Scale bar = 1µm (A), 2µm (D, E).

To further assess and quantify how ER membrane mobility is impacted by desmosome associations, we used a Membrane Displacement Analysis (MDA) macro in Fiji (*16*) to classify ER membrane into static and mobile fractions. If the ER membrane moved >2 pixels (>130nm) between timepoints, we classified it as mobile ER. We drew regions of interest (ROIs) that either encompassed ER tubules proximal to desmosomes (Desmosomal ER) (Fig. 3D) or ER tubules distal to cell-cell contacts (Non-desmosomal ER) (Fig. 3E). Membrane displacement analysis revealed that about 66% of ER membrane present at desmosomes was stable, whereas only 39% of non-desmosomal ER was stable (Fig. 3D-F). These results indicate that ER tubules associated with desmosomes are immobile relative to non-desmosomal ER tubules.

### ER tubules associate with desmosomes during remodeling and assembly

We assessed ER-desmosome interactions as desmosomes underwent remodeling events, such as desmosomal fusion and desmosome assembly (*7, 17*). We observed that ER tubules remain in close association with desmosomes undergoing fusion (Fig. S5A-F). Analysis of overlapping fluorescent signals between desmoplakin-EGFP and mApple-VAPB during desmoplakin puncta fusion revealed that the ER was in contact with at least one desmoplakin puncta prior to fusion (n=67 fusion events analyzed) (Fig. S5G-L). Even after fusion, ER tubules maintained contact with newly fused desmosomes (Fig. S5G-L, bottom row). These ER-DP contacts persisted during successive fusion events, such as three-two-one fusion events (Movie S5). Together, these findings indicate that ER tubules associate with desmosomes under steady-state conditions and during desmosome remodeling events.

To determine the spatiotemporal behavior of ER and desmosomes during *de novo* cell-cell contact formation, we grew cells in low calcium (∼30µM) cell culture conditions for 18-24h to prevent desmosome formation. The calcium concentration was then raised to physiological levels (1.8 mM) to initiate formation of cell-cell contact (*18*). Immediately following the switch to normal calcium levels, we performed live-cell time-lapse maging to track ER tubules as desmosome puncta form at cell-cell contacts. We observed that ER tubules extend towards the cell periphery as cells initiate contact (Fig. 4A-E). When two adjacent cells came into contact, peripheral ER tubules formed mirror images at contacts and no longer retracted from the periphery (n=12 pairs of contacting cells) (Fig. 4B; Movie S6). Interestingly, in all 12 instances where we visualized new cell-cell contact formation, nascent desmoplakin puncta became visible precisely where ER tubules had formed mirror images (Fig. 4C, D). As cell contacts matured, additional ER tubules formed mirror image-like arrangements and immobilized at sites of newly forming desmosome puncta (Fig. 4E).

**Fig. 4.**
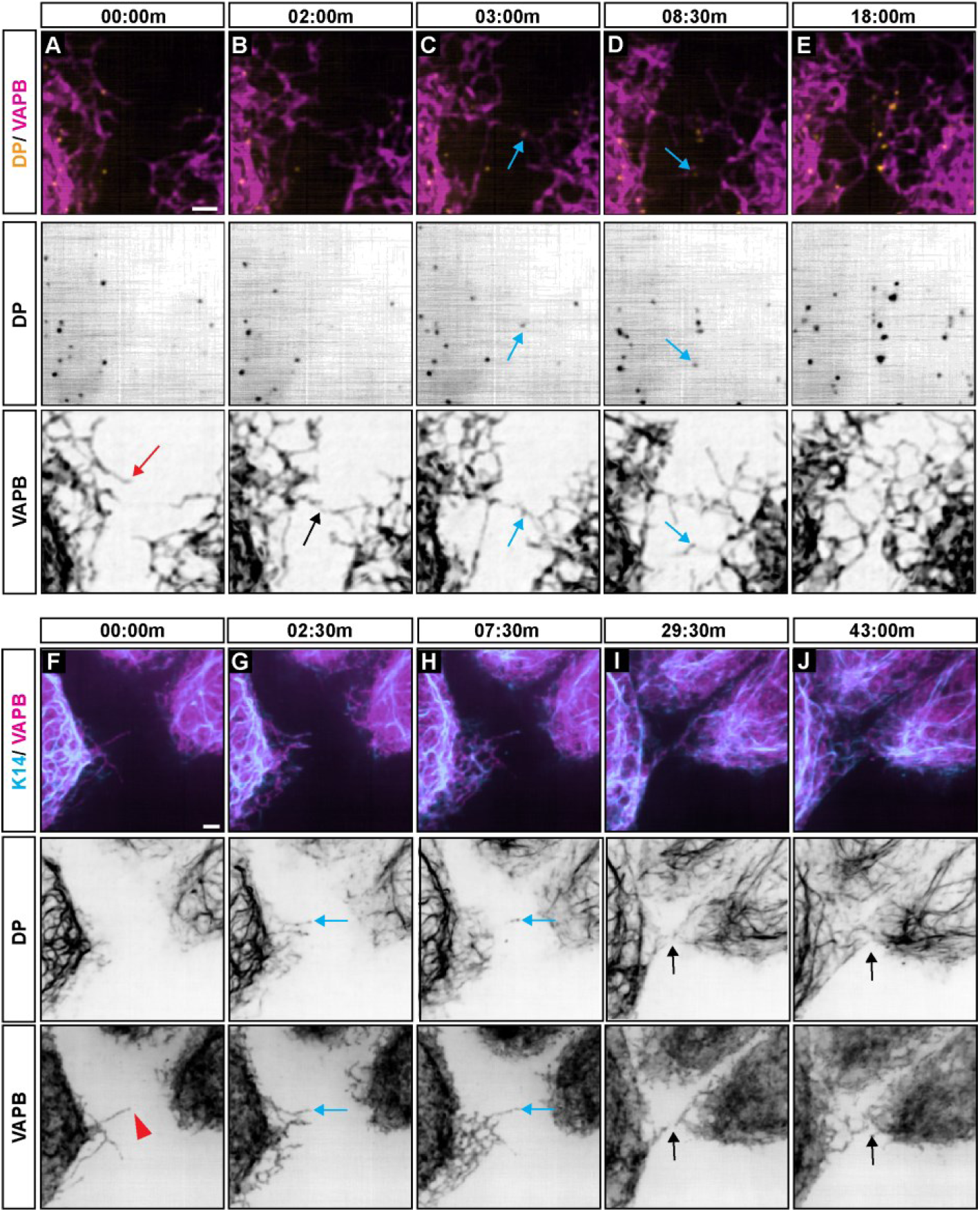
ER tubules associate with desmosomes and keratin filaments during assembly. (**A-E**) Snapshots of a live-cell time-course of desmoplakin (orange) and VAPB (magenta) in A431 cells as they form cell-cell contacts (n=12 samples). ER tubules first extend in both cells (A, red arrow), followed by formation of an ER mirror image at the cell-cell contact (B, black arrow). Desmoplakin puncta appear at the exact position of an ER mirror image (C, D, blue arrows). Eventually, more desmoplakin puncta appear and ER mirror images form as contacts mature (E). (**F-J**) Snapshots of a live-cell time-course of keratin filaments (blue) and VAPB (magenta) in A431 cells as they form cell-cell contacts (n=12 samples). ER tubules sometimes extend alone (F, red arrowhead). ER tubules and keratin filaments extend towards cell-cell contacts simultaneously over several minutes (G, H, blue arrows). Keratin filaments and ER tubules form mirror images as contacts mature (I, J, black arrows). Scale bar = 2µm (A, F).

FIB-SEM imaging revealed a nanoscale association between keratin filaments and ER tubules (Fig. 2). Therefore, we monitored keratin filament and ER tubule dynamics following a calcium switch using A431 cells stably expressing fluorescently tagged keratin-14 (mNeonGreen-KRT14) and mApple-VAPB. Following addition of calcium, some peripheral ER tubules extend and retract without any keratin filaments (Fig. 4F). Eventually, ER tubules and keratin filaments appear to extend to and retract from the periphery simultaneously and in close spatial proximity (Movie S7). These ER-keratin associations persisted for several minutes (Fig. 4G, H). As the cells come into contact, both ER tubules and keratin filaments form stable mirror images on either side of the cell-cell contact (Fig. 4I, J). Collectively, these experiments indicate that desmoplakin puncta formation and keratin filament assembly are initiated at sites of ER tubule extensions at nascent epithelial cell-cell contacts.

### Desmosomes and keratin filaments regulate peripheral ER organization and dynamics

The close contacts and dynamic associations of ER with both desmosomes and keratin filament bundles suggested that the desmosome-keratin adhesive complex might regulate ER organization. To test this possibility, we compared ER and keratin organization in wild type A431 cells and in A431 cells in which the desmosomal cadherin desmoglein-2 (DSG2) gene was ablated using CRISPR/Cas9. We have previously shown that these DSG2-null cells exhibit aberrant localization of desmosomal proteins and weakened cell-cell adhesion strength (*19*). Transmission EM confirms that these cells lack mature desmosomes (data not shown). We stably expressed mNeonGreen-KRT14 and mApple-VAPB in WT and DSG2-null cells and imaged these cells using spinning disk confocal fluorescence microscopy. In WT cells, keratin filaments extend radially towards desmosomal cell-cell contacts. These radial keratin filaments terminate at the cell border where they anchor to desmosomes (Fig. 5A; Movie S8, top row). Peripheral ER tubules exhibit similar organization in these cells, running parallel to the keratin filaments with both structures forming mirror images at cell contacts (Fig. 5A, yellow arrows). In contrast, DSG2-null cells exhibit few radial keratin bundles. Instead, keratins in DSG2-null cells were observed running parallel to the plasma membrane in a subcortical localization (Fig. 5A; Movie S8, bottom row). Similarly, peripheral ER tubules were organized parallel to cell-cell contacts rather than orthogonally to cell-cell borders (Fig. 5A, red arrows). These data indicate that loss of desmosomes alters peripheral ER organization.

**Fig. 5.**
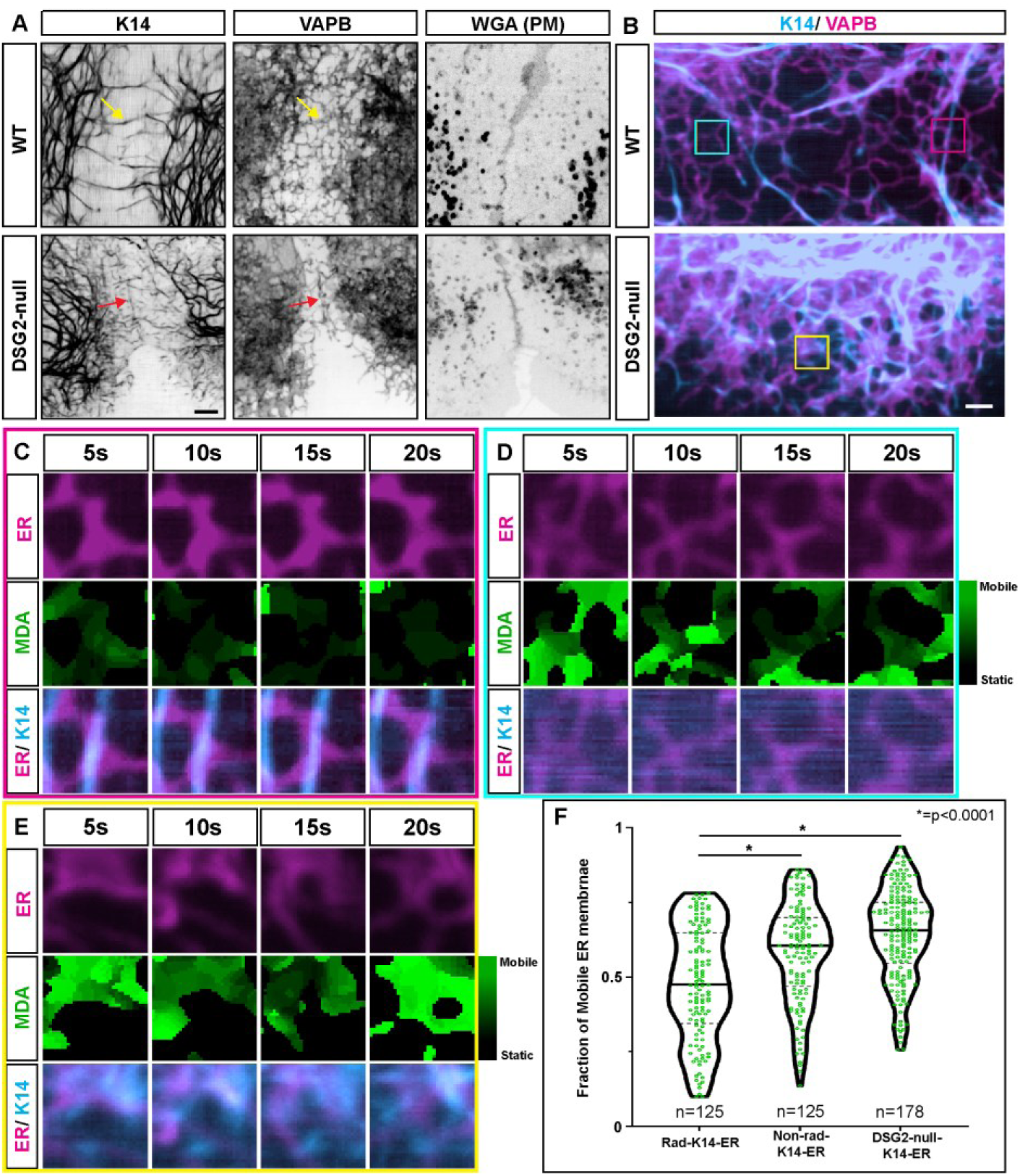
Desmosomes and keratin regulate peripheral ER organization and ER membrane stability. (**A**) A431 WT cells (top row) or Desmoglein2 knockout cells (bottom row) expressing mNeonGreen-KRT14 (first column, keratin marker) and mApple-VAPB (middle column, ER marker). Wheat Germ Agglutinin (WGA) labels the plasma membrane (PM). Yellow arrows show radial keratin filaments and associated VAPB tubules orthogonal to the PM in WT cells. Red arrows show keratin filaments and ER tubules parallel to the PM in DSG2-null cells. (**B**) Light microscopy images of KRT14 (blue) and VAPB (magenta) in A431 WT and DSG2-null cells. Boxes are color-coded and indicate regions analyzed by MDA analysis in C (pink), D (blue), and E (yellow). (**C**) ER (magenta), MDA-generated ER movement (green), ER at radial keratin filaments (merge) in WT cells. (**D**) ER (magenta), MDA-generated ER movement (green), ER at non-radial keratin filaments (merge) in WT cells. (**E**) ER (magenta), MDA-generated ER movement (green), ER at keratin filaments (merge) in DSG2-null cells. Bright green pixels depict ER fraction that is mobile between time points. (**F**) Violin plot depicting fraction of mobile ER. Horizontal black lines in violin plots represent medians (Mann-Whitney, p<0.0001) and *n* indicates number of ROI analyzed. Scale bar = 4µm (A), 2µm (B).

In addition to ER organization, we also assessed the influence of desmosomes on ER dynamics. Membrane displacement analysis was used to determine ER membrane mobility at: 1) ER tubules along radial keratin filaments in WT cells, 2) ER tubules not associated with radial filaments in WT cells, and 3) ER tubules located along keratin filaments in DSG2-null cells (Fig. 5B-E). Analysis revealed that 52% of ER membrane was stable along radial keratin filaments running to desmosomes, whereas only 42% of ER associated with non-radial keratin filaments was stable. These differences were even more stark when assessing ER mobility in DSG2-null cells, with only 36% being stable (Fig. 5F). Together, these results indicate that an association between ER tubules and radial keratin filament bundles suppresses ER tubule dynamics. The stabilization of ER tubules along radial keratin filament bundles explains the persistence of ER tubule positioning in mirror image-like arrangements at desmosomal cell-cell junctions.

### Keratin mutants that cause epidermal blistering alter ER membrane morphology

Keratin mutations cause a wide range of human diseases, often affecting the skin and skin appendages, such as hair and nails (*20*). To determine if disease-causing mutations that disrupt keratin filaments also perturb ER morphology, we stably expressed a KRT14 point mutant, KRT14^R125C^ (mNeonGreen-KRT14^R125C^), in A431 cells expressing mApple-VAPB. This KRT14 mutation acts dominantly to cause aggregation of keratin filaments at the cell periphery, and human patients heterozygous for this mutation present clinically with the skin blistering disease epidermolysis bullosa simplex (EBS) (*21*). Similar to results shown in Fig. 5, KRT14^WT^ formed radial filaments that are closely associated with ER tubules (Fig. 6A-D; Movie S9, top row). In cells expressing the KRT14^R125C^ mutant, peripheral ER morphology was sheet-like or planar, especially when in contact with clusters of keratin aggregates (Fig. 6E-H). The KRT14^R125C^-expressing cells still form some keratin filaments, which might explain why some ER persists as tubules (data not shown). Interestingly, time-lapse spinning disk confocal microscopy over a 2-minute time course revealed that many of the keratin aggregates maintain contact with ER membranes (Movie S9, bottom row). These observations indicate that ER-keratin associations are stable and can occur whether keratin is filamentous or in aggregates (Fig. S6). To verify that changes in ER morphology were not an artifact of differing levels of mApple-VAPB expression in different cells, we plated a mixture of KRT14^WT^ and KRT14^R125C^-expressing A431s (1:1 ratio) onto slides and imaged mixed cell clusters that had similar fluorescence intensities of mApple-VAPB. Again, peripheral ER domains appeared more sheet-like in the KRT14^R125C^-expressing A431s (Fig. 6I-K; Movie S10). These results indicate that a keratin mutation that causes an inherited epidermal blistering disease alters ER membrane morphology.

**Fig. 6.**
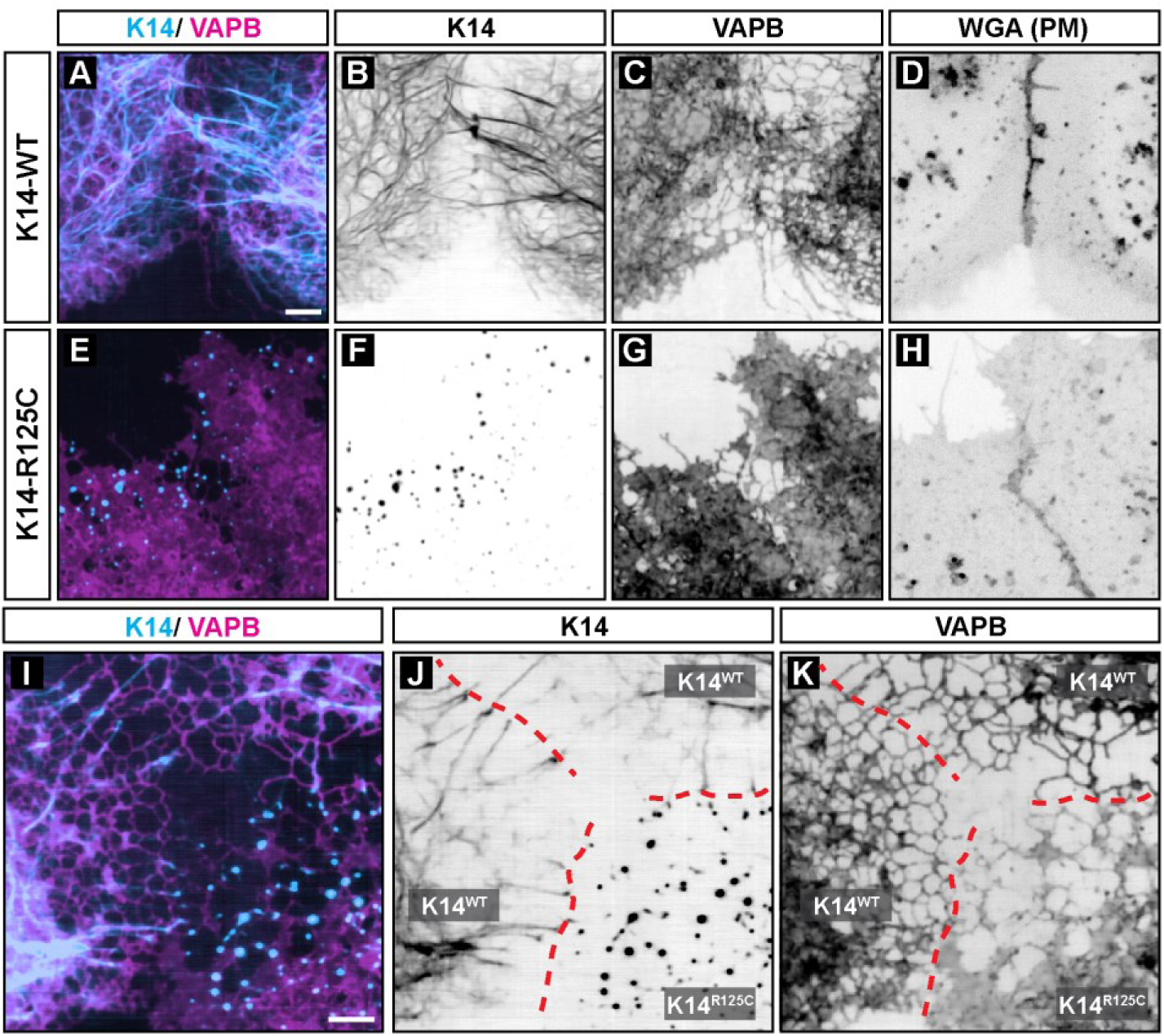
Keratin aggregation perturbs ER membrane morphology. (**A-D**) Light microscopy of K14^WT^, VAPB, and plasma membrane in A431 cells showing ER with tubular morphology. (**E-H**) A431 expressing K14^R125C^ aggregates showing ER with sheet-like morphology. PM is labelled with Wheat Germ Agglutinin (WGA) conjugated with a fluorescent dye. (**I-K**) A431 cells expressing K14^WT^ (top and left cells) or K14^R125C^ (bottom right cell) showing tubular ER in K14^WT^-expressing cells and sheet-like ER membrane in K14^R125C^-expressing cells. Red dashed lines indicate PM. Scale bar = 4µm (A, I).

## Discussion

Our data reveal a previously unknown symmetrical ER junctional organelle formed by the association of three subcellular structures: a desmosome adhesive junction, a keratin intermediate filament scaffold, and peripheral ER tubules. We report that desmosomes regulate peripheral ER morphology by organizing radial keratin filament bundles that position ER tubules orthogonally to the plasma membranes of adjacent cells. These ER tubules exhibit frequent interactions with both keratin filament bundles and the substructure of the desmosome cytoplasmic plaque. This architectural arrangement, along with the characteristic mirror image organization of desmosomes, shapes the morphology and dynamic behavior of peripheral ER membranes at regions of epithelial cell-cell contact.

The ER makes close (<30nm) contacts with the plasma membrane (PM) that function as sites of lipid exchange and calcium homeostasis (*1*). Epithelial cells assemble an extensive array of intercellular contacts that could influence or be regulated by ER-PM associations. A recent study suggested an association between ER and adherens junctions, but neither high-resolution nor dynamic associations between ER and cell-cell junctions were explored (*22*). The ER is associated with highly specialized complexes between Sertoli cells and developing spermatids where it is thought to regulate calcium signaling and junction remodeling (*23*). We find that ER-desmosome associations are stable and occur during desmosome fusion and during initial stages of desmosome formation. When cells are switched from low to high calcium levels to trigger cadherin dependent cell-cell contact, ER tubules extend to the cell periphery as adjacent cells initiate contact. Desmoplakin puncta appear to coalesce *de novo* at the tips of ER tubular extensions. Thus, the ER appears to pattern specific plasma membrane regions for nascent desmosome formation, and reciprocally, newly forming desmosomes function as anchorage points that stabilize peripheral ER tubules.

FIB-SEM imaging at approximately 4nm isotropic voxel size reveals that the ER membrane lies proximal to the desmosome plaque, often within distances observed at canonical ER-PM contacts (Fig. 1; Fig. S3; Movie S1-S3). In several instances, we observe ER tubules in contact with both the inner and outer dense plaque of the desmosome, and in some instances, tubules even penetrate the space between these electron dense structures adjacent to the plasma membrane (Fig. S3). These observations strongly suggest molecular associations between proteins resident on the ER membrane and within the desmosomal substructure. This notion is supported further by optical imaging of living cells. ER tubules associated with desmosomes exhibit reduced mobility compared to non-desmosomal ER (Fig. 3). Knock out of the desmosomal adhesion molecule DSG2 reduced desmosome formation and increased peripheral ER membrane mobility (Fig. 5). Together, these observations indicate that desmosomes control both the organization and dynamics of peripheral ER membranes.

FIB-SEM imaging also revealed interactions between ER tubules and keratin filaments running orthogonally toward desmosomal contacts at the plasma membrane (Fig. 2). In addition, ER membrane frequently surrounded keratin filament bundles (Fig. 2A-C). Consequently, ER tubules and keratin filaments displayed a coordinated spatial and temporal relationship in live cell systems. When initiation of cell-cell contact is triggered with a low-to-high calcium switch, keratin filaments extend along ER tubular extensions, resulting in a stabilized mirror image of tubules and keratin filaments terminating at sites of desmosome maturation (Fig. 4F-J; Fig. 5).

Desmoglein-null cells exhibit a loss of keratin bundles extending out to the cell periphery and a loss of orthogonally organized ER at cell-cell contacts (Fig. 5). Interestingly, expression of a KRT14 mutant that functions dominantly to cause keratin filament aggregation and leads to the epidermal blistering disorder EBS, caused a dramatic reorganization of ER membranes (Fig. 6). In cells expressing wild type KRT14, peripheral ER was predominantly tubular, whereas expression of the keratin EBS mutant reduced tubular ER in favor of planar ER membrane morphology. These findings indicate that desmosomes function to anchor keratin filament bundles which in turn act as filamentous guides that stabilize ER tubules into symmetrical organelle arrangements at epithelial cell-cell junctions. Disruption of either desmosomal or keratin organization profoundly impacts both ER dynamics and morphology.

The identification of a symmetrical ER assembly at desmosomes provides a foundation for fundamentally new ways to conceive of cellular stress sensing mechanisms. Cell stress signaling is a central function of the ER in the regulation of cellular metabolism in homeostatic and disease states (*24*). Likewise, intermediate filaments provide cells and tissues with resistance to mechanical and chemical stresses, particularly in the heart, skin, and liver (*20, 25, 26*). Interestingly, ER stress has been implicated in epidermal fragility disorders caused by keratin or desmosome dysfunction (*27*). Conversely, loss of function mutations in the ER calcium pump SERCA2 cause desmosome and skin defects in Darier’s disease (*28*). These observations suggest that the symmetrical ER-desmosome-keratin structure identified here represents a multifunctional organelle able to resist mechanical forces and mediate cellular stress signaling during disease states.

## Acknowledgments

The authors are grateful to Drs. D. Lerit (Emory University), K. Green and L. Godsel (Northwestern University), and Thomas Magin (University of Leipzig) for instrument use, reagents, and advice. We thank Nate Sheaffer and Joseph Bednarczyk from Penn State College of Medicine’s Flow Cytometry Core for assistance with cell sorting. This research project was supported in part by the Emory University Integrated Cellular Imaging Core.

## Author contributions

NKB, SNS, APK were involved with project conception.

NKB and APK wrote the manuscript with input from all co-authors.

NKB and WG were involved with experimental design, image acquisition and analysis, writing the methods section, and making figures and movies.

TLC supervised CryoSIM and FIB-SEM workflow.

JSA was involved with Cryo light microscopy image acquisition/processing, FIB-SEM data acquisition.

SK was involved with CryoSIM/ FIB-SEM sample preparation including cell culture/labeling, high-pressure freezing, sample trimming.

CPT, SS, and AVW supervised ER segmentation in FIB-SEM datasets.

SP, ET, and DB were involved with FIB-SEM data pre-processing.

DB and ET organized FIB-SEM data and data attributes.

WP and AP provided manual annotations, evaluations, and proofreading.

DB built the data management infrastructure.

LH and SS developed machine learning algorithms; LH performed network training and predictions.

DA and SS developed refinement and analysis algorithms; DA analyzed data.

JB and SS developed automated CLEM registration algorithms; JB performed automated CLEM registration.

AWV was involved with Transmission EM acquisition in Fig. S1.

APK and SNS were involved with funding acquisition.

SNS and KG reviewed the manuscript.

APK supervised the project.

## Data and materials availability

All plasmid maps, code, and macros are available at (*29*). Raw FIB-SEM datasets are hosted on: https://openorganelle.janelia.org. Supplementary Movies S1-S10 can be accessed at: https://youtube.com/playlist?list=PLRVCHiGokrM5RSn2HAUap7ZJBbr82HdKu. An MTA may be required for transfer of some reagents.

## Supplementary Materials

Materials and Methods

Figs. S1 to S7

Tables S1 to S5

Movies S1 to S10

