## Supplementary Materials for "Architecture and dynamics of a novel desmosome-endoplasmic reticulum organelle"

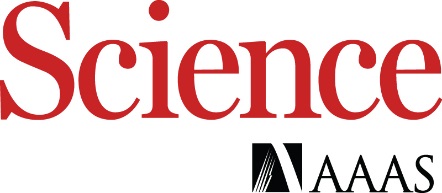

Supplementary Materials for

**Architecture and dynamics of a novel desmosome-endoplasmic reticulum organelle**

Navaneetha Krishnan Bharathan^1^, William Giang^1^, Jesse S. Aaron^2^, Satya Khuon^2^, Teng-Leong Chew^2^, Stephan Preibisch^2^, Eric T. Trautman^2^, Larissa Heinrich^2^, John Bogovic^2^, Davis Bennett^2^, David Ackerman^2^, Woohyun Park^2^, Alyson Petruncio^2^, Aubrey V. Weigel^2^, Stephan Saalfeld^2^, COSEM Project Team^2^, A. Wayne Vogl^3^, Sara N. Stahley^1^, Andrew P. Kowalczyk^1*^

**This PDF file includes:**

Materials and Methods

Figs. S1 to S7

Tables S1 to S5

Captions for Movies S1 to S10

**Other Supplementary Materials for this manuscript include the following:**

Movies S1 to S10

Materials and Methods

**Lentivirus generation**

To generate a lentiviral construct expressing mApple-VAPB, the VAPB ORF (open reading frame) from mCherry-VAPB (*8*) was cloned into the C-terminus of mApple-C1 (*30*) using in vivo assembly (*31*). This step generates mApple-VAPB with a 15bp linker (5’-TCCGGACTCAGATCT-3') in between the mApple and VAPB ORFs. A second round of in vivo assembly was performed to clone mApple-VAPB into the vector backbone of pLenti6-V5-DEST for lentivirus production. The remaining lentiviruses used in this study were generated by VectorBuilder. Vector IDs are specified in Table S1 and can be used to retrieve detailed information about the vector on vectorbuilder.com. The sequences of plasmids we generated in house were confirmed by sequencing and with restriction enzyme digestion. The sequences of all plasmids obtained from VectorBuilder were validated with restriction enzyme digestion. Lentiviruses were made by co-transfection into human embryonic kidney-293FT cells with pMD2.G (encoding VSV-G) and psPAX2 (encoding Gag and Pol) and collection of culture supernatants 24‒72 hours after transfection. Lentivirus was concentrated either by high-speed centrifugation at 4°C for 1.5h at 25,000 rpm, or with the Lenti-X™ Concentrator kit (631231; Takara Bio) following the manufacturer’s protocol. The mCherry-VAPB (gift from Gerry Hammond, Plasmid #108126), mApple-C1 (gift from Michael Davidson, Plasmid #54631), pMD2.G and psPAX2 (gifts from Didier Trono, Plasmid #12259 and #12260, respectively) plasmids were obtained from Addgene. Plasmid sequences are available at (*29*).

**Cell line generation, transfection, culture, and reagents**

A431 cells were cultured in DMEM (Dulbecco’s Modified Eagle Medium) (10-013-CV; Corning) with 10% fetal bovine serum (TET Tested, S10350; R&D Systems) and 1X antibiotic-antimycotic solution (30-004-CI; Corning). CRISPR/cas9 was used to knockout DSG2 (Desmoglein 2) (DSG2 gRNA target sequence GTTACGCTTTGGATGCAAG) as previously described (*19*, *32*). A431 cells stably expressing Tet-On Desmoplakin (DP)-EGFP were generated as described previously (*7*). DP-EGFP expression was induced by culturing the cell lines in 4μg/mL Doxycycline (D3447; Sigma-Aldrich) for 18-24h. A431 cells were transduced with lentivirus by incubating cells with 8-10µg/mL polybrene (TR-1003-G; EMD Millipore) in cell culture media for 24-48h. Cells stably infected with lentiviruses were selected using either blasticidin (2.5-5 μg/mL) (R21001; ThermoFisher Scientific) or puromycin (0.5 μg/mL) (ABT-440; Boston BioProducts) or a combination of both, depending on the lentivirus used. Bulk sorting of cell lines expressing the various constructs was performed by fluorescence activated cell sorting to obtain populations with similar expression levels.

Cells were also transiently transfected with mCherry-VAPB and DP-EGFP plasmids (Fig. S1A-C) using the Viromer RED transfection reagent (TT100302; OriGene Technologies). Briefly, cells were grown to 50% confluency. A mix of the Viromer RED transfection reagent, Viromer RED buffer, and plasmid DNA were added to the cells and incubated. Cells were then imaged 24 hours later. The Desmoplakin-GFP (gift from Kathleen Green, Plasmid #32227**)** was obtained from Addgene.

**Transmission Electron Microscopy**

Cells cultured to 100% confluence on 12mm, 0.4µm, polyester Transwells (CLS3460; Corning) were fixed in 1.5% paraformaldehyde (15713; Electron Microscopy Sciences)/1.5% glutaraldehyde (16320; EMS) in 0.1M sodium cacodylate buffer (pH 7.3) (12300; EMS) at room temperature for 30 minutes. The membrane inserts were then cut out with a clean scalpel and incubated in room temperature fixative. Membranes were then washed 3X with 0.1M sodium cacodylate buffer for 10 minutes each. Samples were then post-fixed on ice for 1h with 1% OsO_4_ (19100;EMS) (1:1 mix of 2% OsO_4_ and 0.2M sodium cacodylate buffer) and then washed 3X (10 minutes each) with ddH_2_O. Membranes were stained with 1% aqueous uranyl acetate and washed again 3X (10 minutes each) with ddH_2_O before being dehydrated in an ascending concentrations series of ethanol solutions, treated with two changes of 100% propylene oxide (20401; EMS), and infiltrated with a 1:1 mix of propylene oxide and EMbed 812 resin (14120; EMS) and left overnight. After two 2hr incubations in 100% EMbed 812 resin, the membranes were placed on a silicone platform with cells facing up and embedding capsules (filled with resin) were inverted onto the membranes. Finally, the samples were heated in a 60°C oven for 48 hours. Polymerized blocks with membranes attached were removed from the silicone platforms and trimmed. Thin sections were cut on a Leica Ultramicrotome and the sections collected on 200 mesh copper grids (G200-Cu; EMS). The sections were stained with uranyl acetate and lead citrate, and imaged on an FEI Tecnai G2 or Talus transmission electron microscope operated at 120 kV.

**Correlative Light and Electron Microscopy (CLEM)**

CLEM was performed according to Hoffman et al., 2020 (*6*) and as described below.

***Sapphire Disk Preparation***

Optically flat sapphire disks (3mm diameter, 50-80 micron thick; Nanjing Co-Energy Optical Crystal Co., Ltd.) were cleaned using basic Piranha solution (5:1:1 mixture of H_2_O : 50% H_2_O_2_ : NH_3_OH) for 4-6 hours at 80-90°C. Disks were then coated (Desk II sputter coater; Denton Vacuum Inc.) with a gold pattern around the outer 0.5mm edge to identify the direction the disk is facing. Disks were then placed in a 20mm MatTek disk, gold side facing down, submerged in 70% ethanol for 2-3 minutes. Ethanol was discarded and disks rinsed 3X in sterile water and let dry. Disks were then coated with 0.1% gelatin in 1:1 H_2_O: growth media for 1 hour at 37°C and left to dry.

***Cell Culture and Staining for CLEM***

Cultured A431 cells stably expressing Desmoplakin-EGFP and mApple-VAPB at 70-80% confluency (maintained at 37°C/5% CO_2_ in 100mm tissue culture dishes) were trypsinized (0.25% Trypsin-EDTA) at 37°C, resuspended and re-cultured onto the non-gold bearing surface of the sapphire disks. After 24hrs, cells were exposed to 4µg/mL Doxycycline for DP-EGFP induction. After 24-48 hours, cells were stained with MitoTracker Deep Red FM (M22426; Molecular Probes; working conc. 25-500nM) at 50nM in a CO_2_ incubator for 15 minutes and washed.

***High-Pressure Freezing***

Sapphire disks containing labeled cells were dipped 3X in freezing media containing Fluorobrite media (A1896702; ThermoFisher), 25% Dextran (Mr ~40,000, 31389-100G; Sigma), and 0.8pM TetraSpeck microspheres (0.2µm diameter, T7280; Invitrogen). They were then placed between aluminum planchettes and subjected to High Pressure Freezing (HPF Compact 01; Wohlwend GmbH) according to manufacturer’s instructions. Samples were then stored in liquid N_2_.

***Optical Imaging***

Specimen fluorescence was imaged using the setup described previously (*6*). Briefly, residual ice on the non-cell bearing side of the sapphire disks was removed via scalpel scraping under liquid N_2_. Samples were then loaded into a custom-built cryostat with an imaging window that maintained sample temperature of ca. 77K for the duration of imaging. Cells were excited via 488nm (4 W, Genesis CX STM; Coherent), 561nm (5 W, 2RU-VFLP-5000-560-B1R; MPB Communications) and 642 nm laser (2 W, 2RU-VFL-P-2000-642-B1R; MPB Communications) illumination and imaged via a 100x, 0.85NA objective lens (CFI L Plan EPI CRB; Nikon) onto an sCMOS camera detector (Orca Flash 4.0 v3; Hamamatsu Corp.).

After correction-collar optimization (using TetraSpeck beads), montage epi-fluorescence images are taken of each coverslip to identify regions of interest (ROIs). Following this, ROIs were imaged using 3D structured illumination microscopy (3D-SIM), with typical field of view (FOV) of 130x130 µm (xy) and 8 µm (z-depth). Images were processed using the SIM reconstruction algorithm reported previously (*33*), with the following typical reconstruction parameters: 0.007 Wiener Filter, 0.7 gamma apodization, 15-pixel radii of the singularity suppression at the OTF origins. Finally, chromatic shifts between each color channel were digitally corrected using the TetraSpeck beads as alignment fiducials.

***Electron Microscopy Preparation***

Samples were unloaded from the Cryo Fluorescence Microscope and again stored in liquid N_2_. They were then transferred to cryotubes containing 1-2% OsO_4_, 0.1% Uranyl Acetate, and 3% water in acetone under liquid N_2_. Freeze substitution was then performed, and sapphire disks embedded in Eponate 12 using the recipe reported previously (*6*). Following these manipulations, Epon resin was removed from the bottom side of the sapphire disk, and the disk removed from the resin block by repeated alternating immersion in warm water and liquid N_2_. The newly exposed Epon surface containing cells was then re-embedded in Durcupan resin.

***X-ray Correlation and Trimming***

Samples were then imaged using a micro X-ray CT system (XRadia 510, Carl Zeiss X-ray Microscopy, Inc.) to identify the positions of the cells of interest. Fluorescence images taken previously were overlayed onto X-ray images to ensure proper ROI location. Specimens were re-mounted on a copper stud and trimmed using an ultramicrotome (EM UC7, Leica Microsystems), with supplementary X-ray CT imaging to guide the trimming process, until the region to be imaged in EM was contained in a resin “tab” with typical dimensions of ca. 100x100x65 microns. Finally, samples were sputter-coated with 10nm gold and 100nm carbon (PECS 682; Gatan) to maintain sample conductivity for EM.

***Focused Ion Beam Scanning Electron Microscopy (FIB-SEM)***

Samples were loaded onto a custom FIB-SEM, consisting of a Zeiss Gemini 450 Field Emission SEM and a Zeiss Capella FIB column oriented at 90 degrees to the SEM beam (*34*). Samples were subjected to repeated FIB milling and SEM imaging to acquire 3D EM datasets. Images taken at 8x8x8nm pixel resolution were imaged at 500kHz readout rate, with 1.2kV landing energy at 2.0nA electron dose. Milling was performed using a 15nA gallium ion beam source at 30kV. Images taken at 4x4x4nm pixel resolution were imaged similarly but with 200kHz readout rate, 0.25nA electron dose, and 0.9KV landing energy. A pipeline based upon render web services (available at: github.com/saalfeldlab/render) was used to align and reconstruct the FIB-SEM images into 3D volumes for analysis. Point match correspondences were extracted using SIFT (*35*), and global optimization (*36*, *37*) was employed to compute per-image affine transformations regularized with a rigid model for each entire volume. After alignment, the volumes were flattened using a spline along the z dimension based on key points that were interactively set in BigDataViewer (*38*).

**Spinning Disk Confocal Microscopy**

***Microscope***

Fluorescence imaging was performed using a Nikon Ti2-E equipped with a Yokogawa CSU-X1 spinning disk unit, LUNF XL laser unit, Nikon Perfect Focus System, Z piezo stage, motorized XY stage, two sCMOS cameras (ORCA-Fusion BT, Hamamatsu Corp.), and two fast filter wheels with most elements controlled through hardware-triggering through the Nikon’s National Instruments Breakout Box. The acquisition was controlled with NIS-Elements v5.30.02 and v5.30.04. The polychroic mirror within the Yokogawa CSU-X1 unit is a Semrock Di01-T405/488/568/647. Single emission filters (Chroma ET525/36m and Chroma ET605/52m) were used with the 488nm and 561nm lasers.

***Live-cell Imaging***

For live-cell imaging, cells were seeded on either 8-well chambered #1.5H cover glass (#C8-1.5H-N; Cellvis) or 35mm #1.5 glass bottom dishes with 4 compartments (#627870; Greiner Bio-One). We used a Nikon 100x/1.49 NA Apo TIRF oil immersion objective with its correction collar optimized for our imaging at 37°C through a #1.5H glass coverslip. Images were taken in 12-bit with high gain (“12-bit Sensitive”) and with “Standard” readout mode. Rough alignment for simultaneous dual-channel imaging was accomplished using 0.1µm multi-color TetraSpeck beads to align the Cairn TwinCam unit (holding a Semrock Di02-R561 beamsplitter) until pixel-perfect overlap was achieved in the center of FOV for the two cameras. Temperature and CO_2_ control for maintaining physiological conditions were provided by a Tokai Hit stage top incubation system (Model STXF-WELSX-SET). Acquisition settings for figures/experiments are listed in Table S2. To accommodate the addition of high calcium media during calcium switch experiments, holes were made in polystyrene lids using heated syringes and then sterilized with 70% ethanol.

**Image processing**

***Automatic ER and mitochondria segmentation in FIB-SEM datasets***

Segmentations for ER and mitochondria were generated using the 8nm and 4nm many-type networks described in (*39*) for DSM-2 and DSM-3, respectively. With a manual validation Fiji plugin (available at: https://github.com/janelia-cosem/Fiji_COSEM_Predictions_Evaluation), we found the optimal network iterations to be 800k for ER, 875k for mitochondria in DSM-2 (from range 800k-1000k) and 625k for ER, 750k for mitochondria in DSM-3 (from range 600k-800k). Voxels predicted to belong to organelles were used to segment the organelles into individual connected components (*39*). Components smaller than 20E6 nm^3^ were removed. In the case of mitochondria, predictions were first smoothed by a Gaussian kernel (σ = 12 nm) prior to connected component analysis.

***FIB-SEM image processing/analysis (after COSEM predictions)***

Predicted organelle segmentation results and raw EM data from COSEM/AIC were cropped to remove regions without biologically relevant voxels. Datasets were then resliced (rotated) to an orientation akin to typical light microscope z-stacks. Keratin intermediate filaments and desmosome outer dense plaque were segmented using ilastik’s pixel classification. The structure and strong electron density of the outer dense plaque permitted predictions from ilastik to be filtered based on size and intensity. Microscopy Image Browser (MIB) (*40*) was used for proofreading segmentation results of keratin filaments and other objects.

Segmentation results, exported from MIB, were imported into Dragonfly along with the corresponding cropped FIB-SEM dataset. Connected component analysis (26-connection) was done to generate individual desmosome objects and to split ER by cell. Sample tearing (from the resin embedding step) led to separate ER objects within the same cell, but these objects were manually merged to be counted as one ER object per cell. Unsigned distance transforms were calculated for both ER objects. The statistics for the minimum intensity of the unsigned distance transform maps for each desmosome object were exported as comma separated value files. Desmosomes at locations of sample tearing were excluded from analysis but not visualization.

To reduce file sizes, the 4x4x4 nm dataset (DSM-3) was chopped into twelve tiles of equal dimensions from which two tiles were selected for segmentation touch-up. More details about FIB-SEM datasets can be found in Table S3. Three-dimensional models in Fig. 1, Fig. 2, Fig. S2, and Fig. S3 were generated using Dragonfly software, Version 2020.2 for Windows (*41*).

***Fluorescence microscopy image processing***

Images were split by channels and timepoints before denoising with either Noise2Void (*42*) or 3DRCAN (Fig. S7) (*43*). After re-combining to a hyperstack, all datasets were corrected for lateral chromatic aberration using NanoJ’s channel registration (*44*). If needed, images were then drift corrected using NanoJ’s drift correction. Macros are provided at (*29*).

***3DRCAN***

A431 cells stably expressing mApple-VAPB were grown on #1.5 glass coverslips and fixed using 2% PFA for 15 minutes at 37°C before being mounted in ProLong Gold and left to cure for 72 hours. Acquisition settings for the low signal-to-noise (SNR) images were chosen to match the SNR during live-cell imaging. The high SNR settings used higher laser power and higher exposure to compensate for the fixation-induced loss in fluorescence. “Ground truth” datasets were obtained after Richardson-Lucy deconvolution of the high SNR datasets. After excluding datasets that showed focal drift between the low/high SNR datasets, we had 37 (34 training and 3 validation) datasets. Training parameters can be found in Table S4. A Windows 10 workstation with an NVIDIA RTX 3090 GPU was used for training/prediction.

***Noise2Void***

Noise2Void v0.2.1 was used with TensorFlow-DirectML on a Windows 10 workstation with an NVIDIA RTX 3090 GPU (Graphics Processing Unit). In some cases, N2V training and prediction used the N2V CSBDeep Fiji plugin on a Windows 10 workstation with an NVIDIA Quadro RTX 4000 GPU. Training parameters can be found in Table S5.

***CLEM Registration***

Fluorescence and EM image data were coregistered as follows. The mitochondria fluorescence channel (MitoTracker) was manually cropped to approximately match the field of view of the FIB-SEM image. Mitochondria predictions (see *Automatic ER and mitochondria segmentation in FIB-SEM datasets* section) were downsampled to 64nm isotropic resolution and coarsely aligned to the cropped light image using BigDataViewer (*38*). Resampled predictions were then blurred with a 1x1x3 pixel Gaussian kernel using Fiji (*45*). Processed predictions were registered to the fluorescence image using elastix (*46*) using two steps. First, we used elastix to estimate an affine transformation between the two volumes. We then ran elastix again, initialized with the estimated affine to find a non-linear transformation. Parameters for both steps can be found at (*29*)<https://github.com/KowalczykLab/Desmosome-ER>. In estimating the nonlinear transformation, we provided elastix an automatically generated mask that was non-zero when the image fields of view overlapped. The inverse of the resulting transformation was estimated and applied to all fluorescence images, bringing them into spatial alignment with the FIB-SEM (using code from: [https://github.com/saalfeldlab/template-building](https://urldefense.com/v3/__https:/github.com/saalfeldlab/template-building__;!!Ls64Rlj6!nlp886iXSm5NsowEN8--MU8XbvdyP0wkKdSozKFU0JW32krXGX27iBjYZcuy74kZ5p8XHGWE8A$)).

**Image Analysis**

***Segmentation with ilastik and Generation of overlapping signals***

To determine the extent of contact between DP and VAPB over time, we performed an analysis of overlapping signals (Fig. 3C). We first generated binary masks of both DP-EGFP and mApple-VAPB channels using the Pixel Classification workflow in ilastik (*47*). Briefly, features were drawn to differentiate foreground from background pixels in denoised images. This step generates a binary image for each channel, with true signal value=1 and background value=0.

To generate an overlap image, we used Fiji (*45*) to perform a “Multiply” calculation on ilastik-segmented binary masks of corresponding DP and VAPB images. Therefore, only pixels that have a true signal (value=1) in both channels will be displayed. These resulting pixels suggest contact between DP (desmosomes) and VAPB (ER) at the resolution limit of standard spinning disk confocal microscopy.

***Analysis of DP-VAPB contact***

Fiji plugins were developed to determine whether DP puncta made contacts with ER membrane over a 2-minute duration with image acquisition every 5 secs (25 timepoints) (Fig. 3C). Briefly, we drew ROIs around individual ilastik-segmented DP puncta to obtain information about puncta position and area. Only DP puncta that remained in focus for all 25 timepoints and that were equal to or larger than 4µm^2^ were selected for analysis. These ROI positions were then applied to the corresponding “Multiplied” image file (see *Segmentation with ilastik and Generation of overlapping signals* section) to acquire area measurements at each timepoint. An area value of 0µm^2^ meant no overlap between a DP ROI and ER membrane for that timepoint.

***Membrane Displacement Analysis (MDA)***

To quantify the displacement of ER membranes at various cellular locations and between conditions, we calculated optical flow fields and classified membrane movement as either static or mobile using a modified version of a previously published Fiji macro (*16*). We performed MDA on ROIs of 35px X 35px (2.275µm X 2.275µm) with the following parameters for MSEGaussianFlow: (sigma=4, maximal_distance=10). The extent of membrane movement was depicted by the pixel intensity, with higher movement between frames corresponding to higher intensity values. ER membrane movement was classified into two categories based on the amount of movement in pixels: “Static ER” (0-2 pixels) or “Mobile ER” (3-9 pixels). Masking of the mApple-VAPB channel was performed using the “Percentile” thresholding in Fiji. The MSEGaussianFlow plugin was authored by Stephan Saalfeld and Pavel Tomancak.

**Statistical analysis**

Statistical testing was performed using Prism 8 (GraphPad Software) as indicated in the figure legends (Fig. 3F and Fig. 5F). Normality testing was first performed with the D'Agostino-Pearson omnibus (K2) test. When normality tests failed, statistical significance was calculated using the nonparametric Mann-Whitney test (p<0.001).

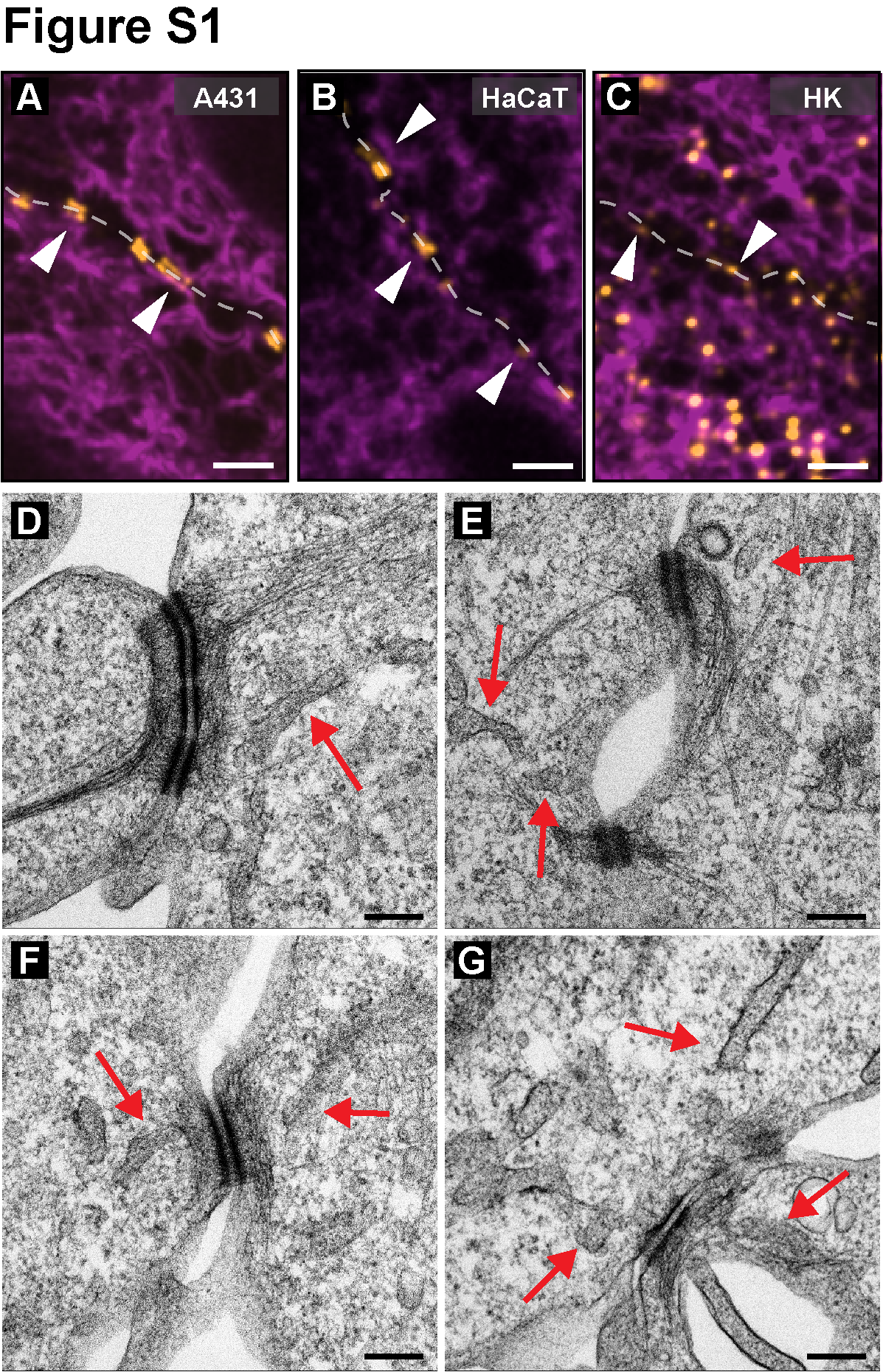

**Fig. S1. ER tubules are proximal to desmosomal junctions.** (**A-C**) Images of a pair of A431 cells (**A**), HaCaT immortalized keratinocytes (**B**), and primary Normal Human Epidermal Keratinocytes (**C**) expressing mCherry-VAPB (magenta, ER marker) showing ER tubules proximal to desmosomal junctions (white arrowheads). Desmosomes are labelled with DP-EGFP in A, B and with an Alexa Fuor-488-conjugated anti-DSG3 monoclonal antibody (AK23) in C. (**D-G**) Transmission electron micrographs in A431 epithelial cells show ER tubules (red arrows) proximal to the electron-dense desmosomal plaque. Scale bar = 2µm (A-C), 200nm (D-G).

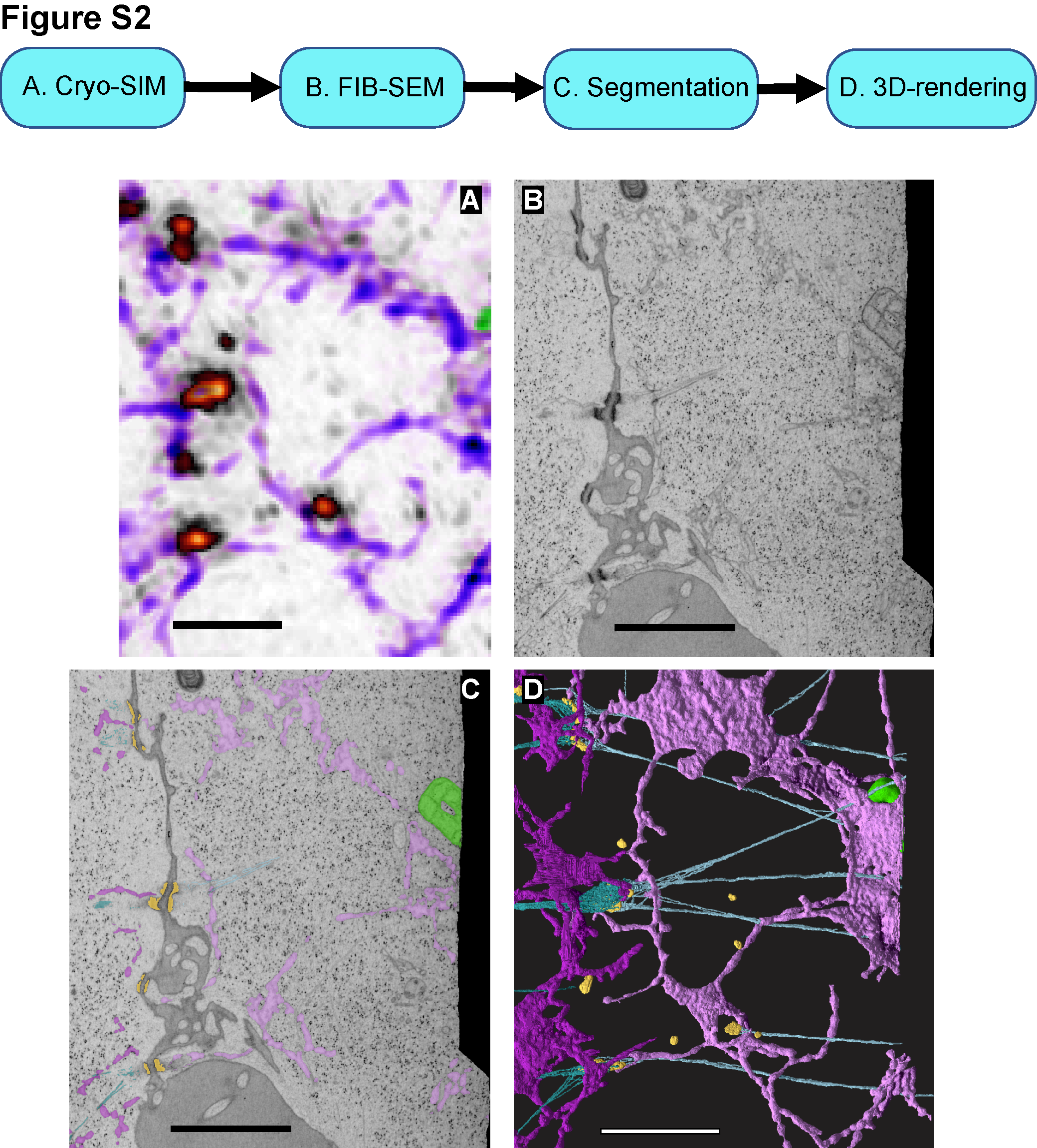

**Fig. S2. Correlative Light and Electron Microscopy (CLEM) workflow.** (**A**) A light microscopy Cro-SIM image of a cell-cell contact with labelled desmosomes (orange), ER (purple), and mitochondria (green). (**B**) A FIB-SEM slice of the same region in A. (**C**) Segmentation of the desmosome outer dense plaque (orange), ER (magenta), and mitochondria (green) in a FIB-SEM slice. (**D**) Three-dimensional reconstruction of the structures segmented as shown in C. Scale bar = 2µm.

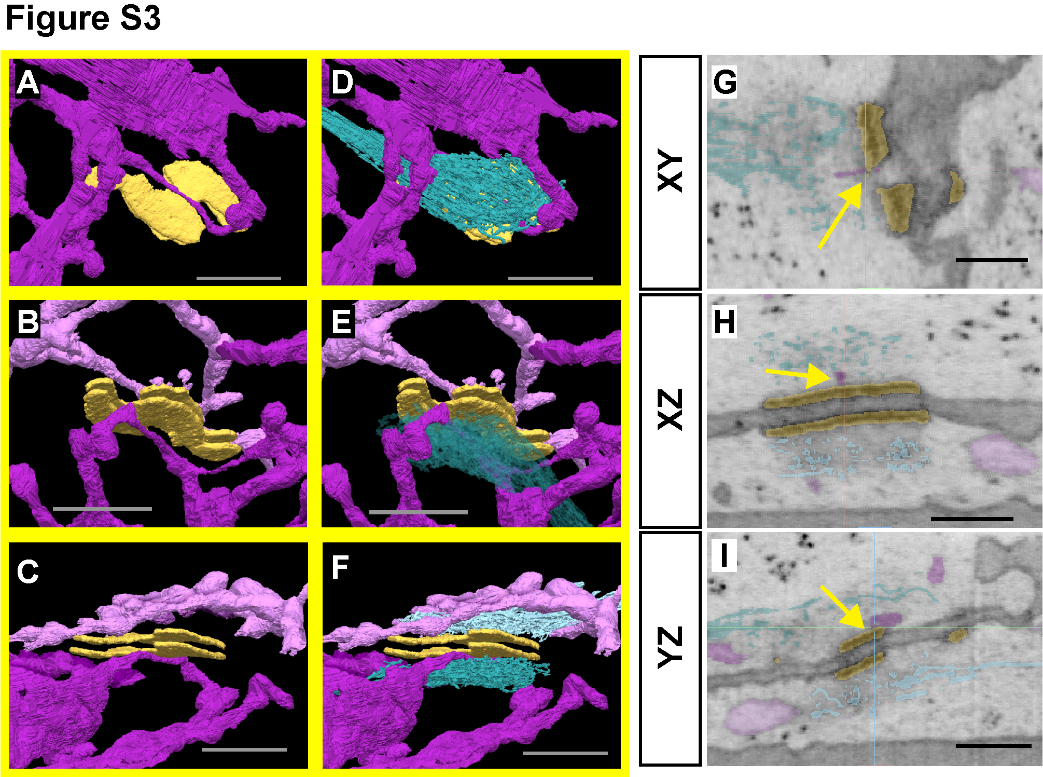

**Fig. S3. FIB-SEM reveals symmetrical keratin and ER tubules on either side of the desmosome contact.** Magnified and rotated views of yellow box in Fig. 1B. (**A-C**) Rotated views showing magenta ER tubules making contact with orange desmosome outer dense plaque. (**D-F**) Rotated views showing teal keratin filaments on either side of the orange desmosome outer dense plaque. (**G-I**) Orthoslices in XY (**G**), XZ (**H**), and YZ (**I**) with desmosome, ER, and keratin segmentations. Yellow arrows point to contact between ER tubules and the desmosome outer dense plaque in all 3 views. Scale bar = 500nm (A-F), 250nm (G-I).

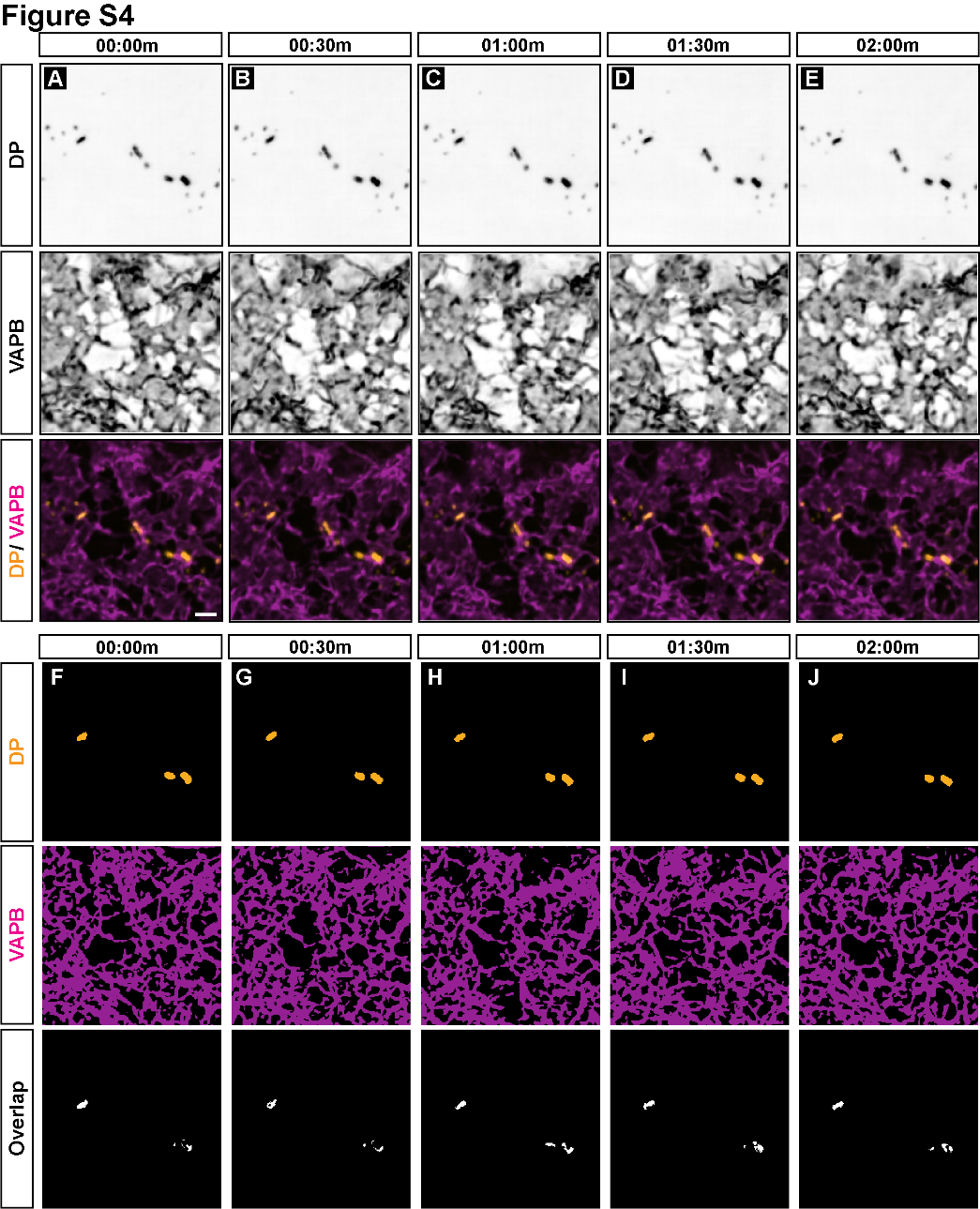

**Fig. S4. ER tubules maintain contact with desmosomes in live cells.** (**A-E**) Snapshots of a live-cell time-course of desmoplakin (top row, orange in bottom row) and VAPB (middle row, magenta in bottom row) in A431 cells at the cell-cell contact over 2 minutes. (**F-J**) ilastik-rendered segmentations of DP (top row) and VAPB (middle row) channels in A-F. Bottom row indicates only overlapping pixels (white) between DP and VAPB channels. Overlap indicates that ER tubules maintain contact with DP puncta. Scale bar = 2µm.

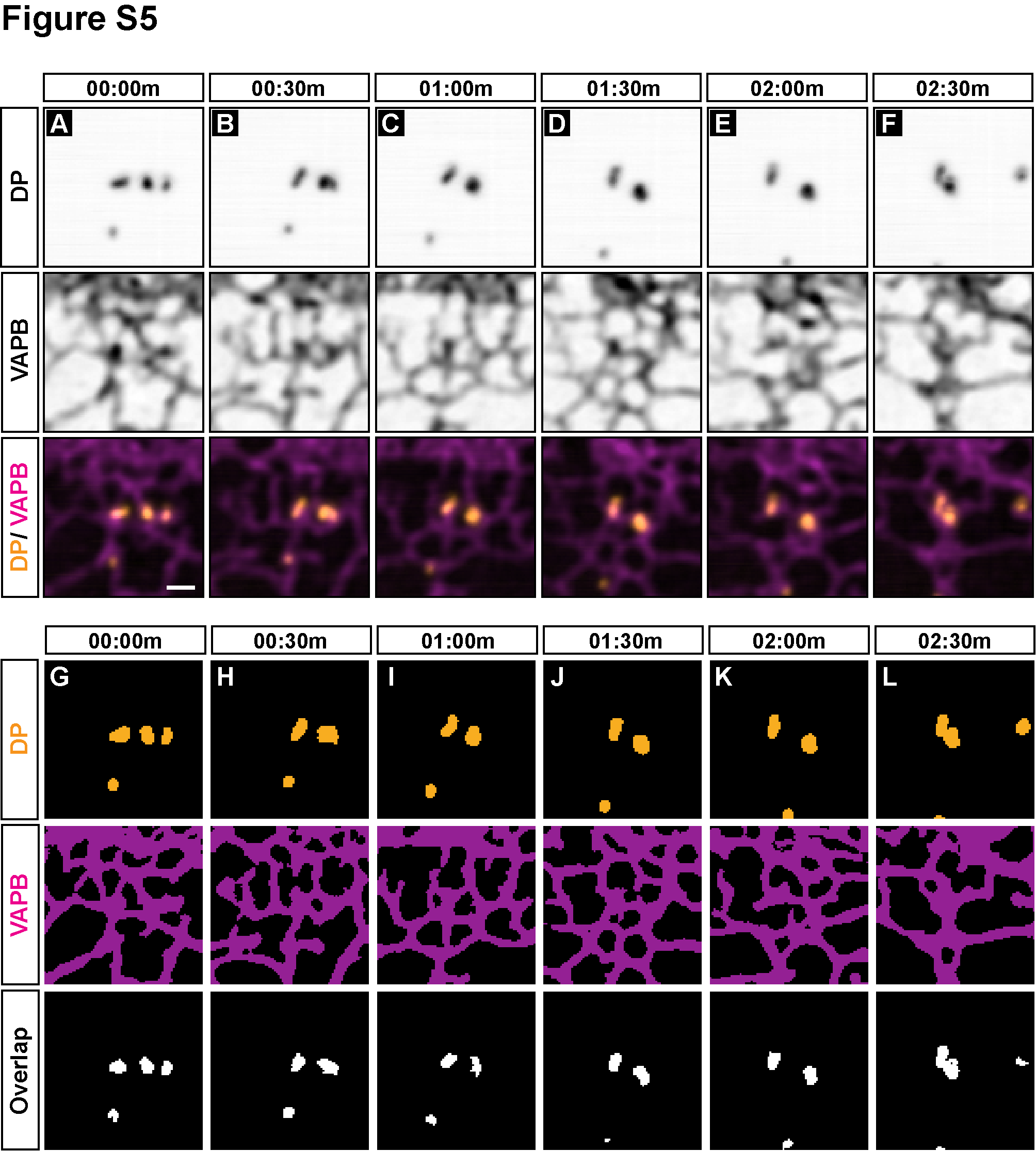

**Fig. S5. ER tubules associate with desmosomes during fusion events.** (**A-F**) Snapshots of a live-cell time-course of desmoplakin (top row, orange in bottom row) and VAPB (middle row, magenta in bottom row) in A431 cells during the fusion of DP puncta at the cell-cell contact. Three DP puncta (A) fuse to form two puncta (B). Eventually, these puncta fuse to form one DP puncta (F). (**G-L**) ilastik-rendered segmentations of DP (top row) and VAPB (middle row) channels in A-F. Bottom row indicates only overlapping pixels (white) between DP and VAPB channels. Overlap indicates that ER tubules maintain contact with DP puncta during fusion. Scale bar = 1µm.

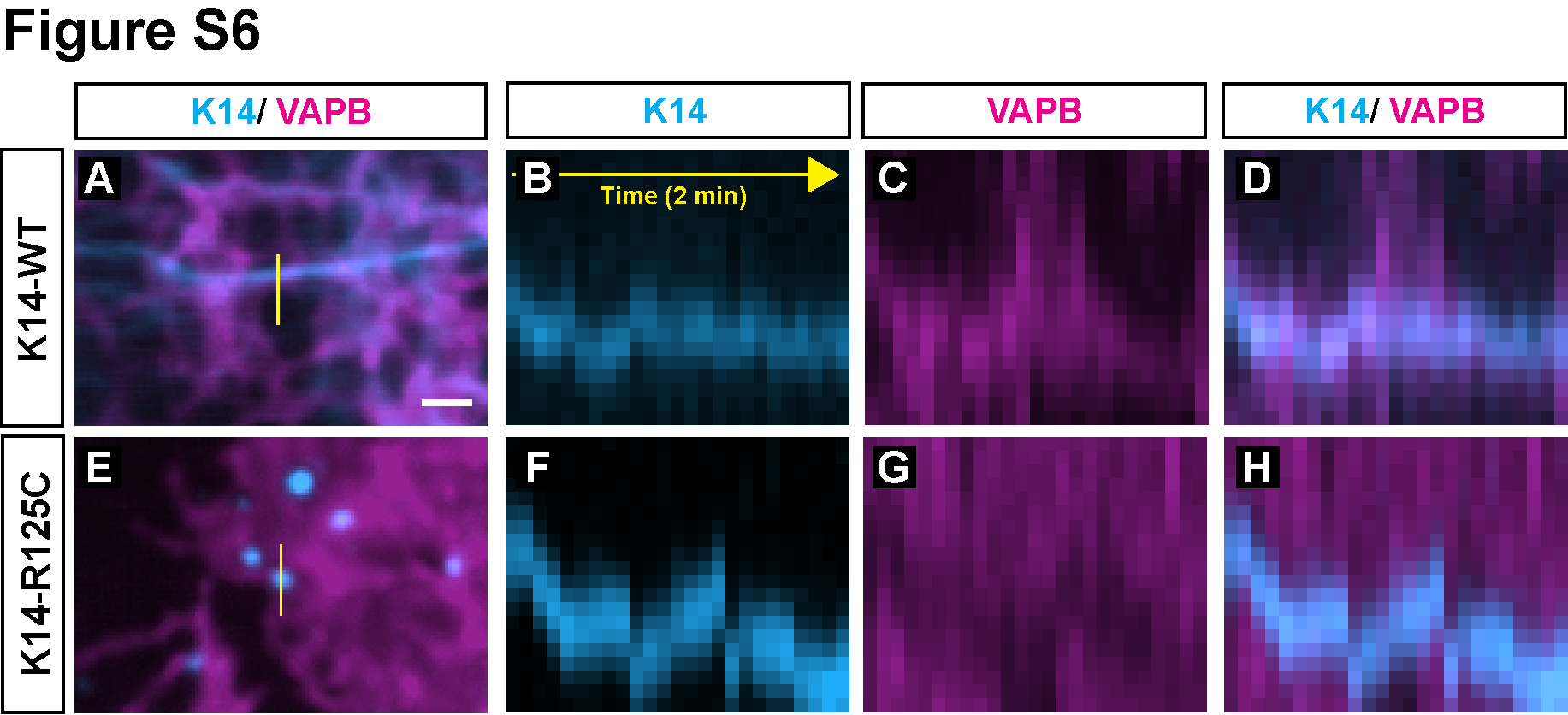

**Fig. S6. Keratin filaments and aggregates are stably tethered to ER membranes.** (**A**) Snapshot of an A431 cell expressing mNeonGreen-KRT14^WT^ (blue) and mApple-VAPB (magenta) showing ER tubules along keratin filaments. Solid yellow line indicates position of kymograph in B-D. (**B-D**) Kymograph of yellow line in A revealing stable ER-keratin contacts over a 2-minute time course. (**E**) Snapshot of an A431 cell expressing mNeonGreen-KRT14^R125C^ (blue) and mApple-VAPB (magenta) showing ER sheets surrounding K14^R125C^ aggregates. Solid yellow line indicates position of kymograph in F-H. (**F-H**) Kymograph of yellow line in E revealing stable ER-keratin contacts over a 2-minute time course. Scale bar = 1µm.

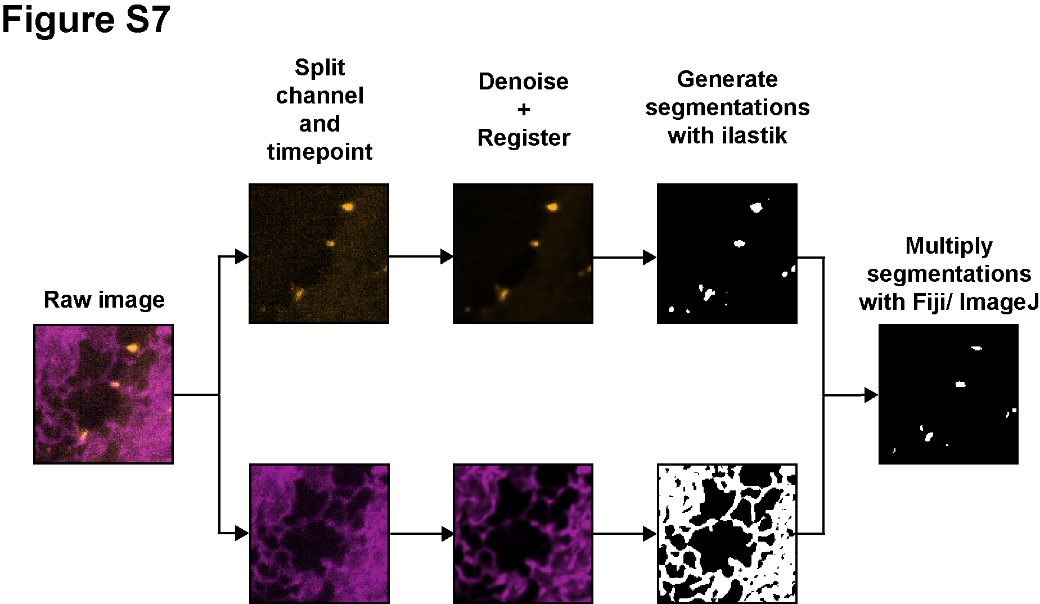

**Fig. S7. Schematic flow of image processing and analysis to determine DP and VAPB contact.** Raw images are first acquired and are then split by channel and timepoint in Fiji/ImageJ. These images are then denoised using 3DRCAN or Noise2Void, followed by registration. Segmentation of channels is performed with ilastik Pixel Classification to obtain binary masks. These image masks are then “Multiplied” in Fiji/ImageJ to detect overlapping pixels, indicating close contact.

| **Vector** | **Insert details** | **Vector ID (VectorBuilder)** | **Antibiotic selection markers** |
| --- | --- | --- | --- |
| pLV-CMV-mNeonGreen-hKRT14 | Expresses wildtype human Keratin 14 (NM_000526.5) tagged with mNeonGreen at the N-terminus. | VB210812-1168rub | Ampicillin/Puromycin |
| pLV-CMV-mNeonGreen-hKRT14-R125C-EBS | Expresses human Keratin 14 (NM_000526.5) containing a c.373C>T substitution and tagged with mNeonGreen at the N-terminus. | VB210812-1169cvr | Ampicillin/Puromycin |

**Table S1.**

Lentiviral vectors used in this study.

| **Figure** | **Experiment** | **Number of slices** | **Z-step size [µm]** | **Exposure time [ms]** | **Interval [s]** | **Duration [min]** |
| --- | --- | --- | --- | --- | --- | --- |
| Fig. 3, Fig. S4 | DP/VAPB fast | 5 | 0.3 | 80 | 5 | 2 |
| Fig. 4A-E | Calcium switch (DP/VAPB) | 7 | 0.45 | 50 | 30 | 90 |
| Fig. S5 | DP/VAPB fusion | 7 | 0.45 | 50 | 120 | 30 |
| Fig. 4F-J | Calcium switch (K14^WT^/VAPB) | 7 | 0.45 | 60 or 70 | 30 | 90-120 |
| Fig. 5, Fig. 6A-D, Fig. S6A-D | K14^WT^/VAPB | 5 | 0.45 | 70 | 5 | 2 |
| Fig. 6E-H, Fig. S6E-H | K14^R125C^/VAPB | 5 | 0.45 | 60 or 70 | 5 | 2 |
| Fig. 6I-K | K14^WT^/VAPB and K14^R125C^/VAPB mixed | 7 | 0.45 | 60 | 5 | 2 |

**Table S2.**

Spinning disk confocal acquisition settings.

|  | **DSM-2 (Fig. 1A)** | **DSM-3 (Fig. 1B-K, Fig. 2, Fig. S3)** |
| --- | --- | --- |
| **Voxel size [nm X nm X nm]** | 8 x 8 x 8 | 4 x 4 x 4 |
| **Total Voxel count** | 5285 x 6566 x 1302 | 3736 x 12101 x 3233 |
| **Total Volume [µm X µm X µm]** | 42.3 x 52.5 x 10.4 | 14.9 x 48.4 x 12.9 |
| **ROI voxel count** | 3217 x 1584 x 447 | 1868 x 2017 x 1001 |
| **ROI volume [µm X µm X µm]** | 25.7 x 12.7 x 3.6 | 7.5 x 8.1 x 4.0 |

**Table S3.**

FIB-SEM datasets analyzed in this study.

| **Figure** | Fig. 4A-E, Fig. S4, Fig. S5 (VAPB channel only) |
| --- | --- |
| **Number of epochs** | 300 |
| **Steps per epoch** | 400 |
| **Number of residual groups** | 5 |
| **Number of residual blocks** | 5 |
| **Patch size** | [8, 128, 128] |
| **Initial learning rate** | 1e-4 |

**Table S4.**

Training parameters for 3DRCAN.

| **Figure** | **Fig. 4A-E, Fig. S4, Fig. S5 (DP)** | **Fig. 3 (DP)** | **Fig. 3, Fig. 4F-J, Fig. 5, Fig. 6, Fig. S6 (VAPB)** | **Fig. 4F-J, Fig. 5, Fig. 6, Fig. S6**  **(KRT14^WT^ and KRT14^R125C^)** |
| --- | --- | --- | --- | --- |
| **Number of epochs** | 400 | 500 | 150 | 150 |
| **Steps per epoch** | 300 | 250 | 200 | 200 |
| **Patch Shape** | 160 | 64 | 64 | 64 |
| **Initial learning rate** | 4e-4 | 4e-4 | 4e-4 | 4e-4 |
| **Batch size** | 80 | 180 | 64 | 64 |
| **Python or Fiji** | Fiji | Fiji | Python | Python |

**Table S5.**

Training parameters for Noise2Void.

**Movie S1. Correlative cryo-SIM/FIB-SEM reveals a novel desmosome-endoplasmic reticulum complex.** FIB-SEM of the cell-cell contacts in A431 epithelial cells shows segmentations of desmosome outer dense plaques (orange) and keratin filaments (teal). Endoplasmic reticulum tubules (magenta) are proximal to desmosome plaques and keratin filaments, forming mirror images on either side of the cell-cell contact. Plasma membrane is labelled a reddish orange.

**Movie S2. FIB-SEM reveals ER-desmosome contact.** Segmentations reveal ER tubules (magenta) in the space between the keratin filaments (teal) and the desmosome outer dense plaque (orange). ER tubules are in contact with the desmosome.

**Movie S3. FIB-SEM reveals ER-keratin association.** Segmentations reveal ER tubules (magenta) wrap around the keratin filaments (teal), making close contact.

**Movie S4. ER tubules form stable mirror images at desmosomal junctions.** Live-cell spinning disk confocal microscopy of A431 epithelial cells shows that ER tubules (magenta) are stably anchored to desmoplakin puncta (orange) over a 2-minute duration forming mirror images (yellow arrowheads) on either side of the cell-cell contact. Scale bar = 2µm.

**Movie S5. ER tubules associate with desmosomes during fusion events.** Live-cell spinning disk confocal microscopy of A431 epithelial cells shows that ER tubules (magenta) persist at desmoplakin puncta (orange) as puncta undergo fusion (yellow box in left panel, red box in middle and right panel). Scale bar = 4µm.

**Movie S6. ER tubules** **associate with desmosomes during cell-cell contact formation.** Live-cell spinning disk confocal microscopy of A431 epithelial cells shows ER tubules (magenta) forming mirror images at sites of nascent desmoplakin puncta formation (orange) (yellow arrowhead in left panel, black arrowheads in middle and right panel). Scale bar = 2µm.

**Movie S7. ER tubules associate with keratin filaments during cell-cell contact formation.** Live-cell spinning disk confocal microscopy of A431 epithelial cells shows ER tubules (magenta) and keratin filaments (blue) extending together as cells form contacts (at 2.5 min.). Eventually, at 29.5 min., ER tubules and keratin filaments form mirror images at cell-cell contacts (yellow arrowhead in left panel, black arrowheads in middle and right panel). Scale bar = 2µm.

**Movie S8. Desmosomes regulate peripheral ER tubule organization.** Live-cell spinning disk confocal microscopy of a pair of A431 WT cells (top row) or A431 DSG2-null cells (bottom row). In A431 WT cells, keratin filaments (blue) and ER tubules (magenta) extend radially towards the cell-cell contact (yellow arrowheads). In A431 DSG2-null cells, keratin filaments (blue) and ER tubules (magenta) are perpendicular to the cell-cell contact (red arrowheads). Scale bar = 4µm.

**Movie S9. Keratin filaments regulate peripheral ER morphology.** Live-cell spinning disk confocal microscopy of a pair of A431 cells expressing KRT14^WT^ (top row) or the EBS (Epidermolysis Bullosa Simplex) mutant KRT14^R125C^ (bottom row). In KRT14^WT^ cells, keratins form filaments (blue) and peripheral ER morphology is tubular in nature. In KRT14^R125C^ cells, keratins form aggregates (blue) and peripheral ER morphology is planar/sheet-like in nature. Scale bar = 4µm.

**Movie S10. ER membranes stably associate with keratin filaments or aggregates.** Live-cell spinning disk confocal microscopy of a mix of A431 cells expressing KRT14^WT^ (top and left cells) or the EBS mutant KRT14^R125C^ (bottom right cell). In KRT14^WT^ cells, keratins form filaments (blue) and stably associate with peripheral ER tubules (yellow box). In KRT14^R125C^ cells, keratins form aggregates (blue) and stably associate with planar ER membranes (red box). Scale bar = 4µm.
